# Mg^2+^ Impacts the Twister Ribozyme through Push-Pull Stabilization of Non-Sequential Phosphate Pairs

**DOI:** 10.1101/742940

**Authors:** A. A. Kognole, A. D. MacKerell

## Abstract

RNA molecules perform a variety of biological functions for which the correct three-dimensional structure is essential, including as ribozymes where they catalyze chemical reactions. Metal ions, especially Mg^2+^, neutralize these negatively charged nucleic acids and specifically stabilize RNA tertiary structures as well as impact the folding landscape of RNAs as they assume their tertiary structures. Specific binding sites of Mg^2+^ in folded conformations of RNA have been studied extensively, however, the full range of interactions of the ion with compact intermediates and unfolded states of RNA is challenging to investigate and the atomic details of the mechanism by which the ion facilitates tertiary structure formation is not fully known. Here, umbrella sampling combined with oscillating chemical potential Grand Canonical Monte Carlo/Molecular Dynamics (GCMC/MD) simulations are used to capture the energetics and atomic-level details of Mg^2+^-RNA interactions that occur along an unfolding pathway of the Twister ribozyme. The free energy profiles reveal stabilization of partially unfolded states by Mg^2+^, as observed in unfolding experiments, with this stabilization being due to increased sampling of simultaneous interactions of Mg^2+^ with two or more non-sequential phosphate groups. Notably, the present results indicate a push-pull mechanism where the Mg^2+^-RNA interactions actually lead to destabilization of specific non-sequential phosphate-phosphate interactions while other interactions are stabilized, a balance that facilitates the folding and stabilization of Twister including formation hydrogen bonds associated with the tertiary structure. The present study establishes a better understanding of how Mg^2+^ ion-interactions contribute to RNA structural properties and stability.

**Statement of Significance:** RNAs are biologically and therapeutically of great emerging interest such that it is critical to understand how RNA molecules fold into complex structures. While experiments yield information on the stabilization of RNA by ions they are limited in the atomic-level insights they can provide. A combination of enhanced sampling methods is applied to explore the compact intermediate states of RNA and their interactions with Mg^2+^ ions. Results reveal a picture of how Mg^2+^ overall stabilizes short phosphate-phosphate interactions thereby facilitating the stabilization of RNA, though doing so by both the stabilization and destabilization of specific interactions. The applied method will be applicable to exploring the impact of divalent ions on the conformational heterogeneity of a range of macromolecules.

## Introduction

RNA molecules have highly diverse structures ranging from simple helices to highly heterogeneous folded conformations that are essential for their wide range of cellular functions (1-3). Specifically, ribozymes, a distinct class of enzymes, exhibit complex tertiary structures and catalyze self-cleavage or the cleavage of phosphodiester bonds of substrate RNA, with metal ions typically playing a central role in the catalytic activity (4, 5). To assume their tertiary structures RNAs must overcome large unfavorable electrostatic interactions associated with their polyanionic phosphodiester backbone (6). To facilitate this, positively charged ions screen the highly negative potential allowing the RNA secondary structures to collapse into compact tertiary conformations (7-10). Typically divalent ions, most often Mg^2+^, facilitate the folding of RNA into tertiary interactions (11-13). However, the inability to visualize the ions during folding represents a key barrier to understanding the role of divalent ions in folding of RNA (14).

Studies have used classical MD or other theoretical approaches to investigate Mg^2+^-RNA binding, but they were limited to native conformations due to their inability to overcome the issues associated with the Mg^2+^ exchange rates (15-21). The exchange rate of water complexed to Mg^2+^ is on the µs time scale (6.7 × 10^5^ s^-1^) and the exchange rate of Mg^2+^ with phosphate is on the ms time scale (0.5 to 2.5 x 10^3^ s^-1^) (22), which is beyond the time scale of typical atomistic MD simulations (19, 23) such that only limited insights into Mg^2+^-RNA interactions are accessible (24). Alternatively, simulations using coarse-grained models of nucleic acids provided insights into how Mg^2+^ can serve to nucleate the folding of key tertiary interactions with the Mg^2+^-RNA interactions being dominated by specific interactions even in unfolded states (18, 25). Reduced models were able to reproduce thermodynamics of Mg^2+^-RNA interactions but were limited to native states (26). Atomistic simulations of unfolding of a pseudoknot at high temperatures showed diverse intermediate states, although divalent ions were absent in those studies (27). Additionally, scientists have worked on improving the force-field parameters for positively charged metal ions to better simulate their interactions with biomolecules (28-30). Overall, a detailed picture of the interactions of Mg^2+^ in an explicit solvent environment with unfolded states of RNA has not yet been attained.

To investigate the impact of Mg^2+^ on the folding and stabilization of RNA at an atomic level of details we apply umbrella sampling MD simulations in conjunction with oscillating chemical potential (μ_ex_) Grand Canonical Monte Carlo (GCMC) sampling (31). The application of GCMC allows for redistribution of Mg^2+^ ions thereby addressing the issue of exchange rates while the MD allows the RNA and waters to respond to the changes in ion positions along a folding pathway. The use of GCMC in combination with MD on RNA was first undertaken by Lemkul et. al. on four structurally distinct RNAs in their native conformations with restraints on the backbone and was shown to successfully identify experimental Mg^2+^ binding sites as well as predict new ion binding regions (32). In the present study we extend that approach by combining it with umbrella sampling to sample the distribution of Mg^2+^ around intermediate conformations along a folding pathway. Application of the method reveals an atomic picture of how Mg^2+^ lowers the free energy of partially folded states of RNA by increasing the sampling of specific non-sequential phosphate-phosphate interactions through simultaneous indirect interaction between two or more non-sequential phosphates. At the same time, other non-sequential phosphate-phosphate interactions are shown to be destabilized as required to allow for overall stabilization of the phosphate-phosphate interactions, representing a push-pull mechanism by which Mg^2+^ stabilizes RNA.

The Twister ribozyme was selected for the present study based on the availability of a range of structural and biochemical data (33-38). While Twister sequences are extremely widespread, the crystal structure (PDB 4OJI) of the ribozyme used in this study is based on Osa-1-4 sequence from *Oryza Sativa* (34). Figure 1 illustrates the secondary and tertiary interactions in Twister. The studied ribozyme has the conserved residues in loops L1, L2 and L4 corresponding to the self-cleavage site and major tertiary interactions (T1 and T2) associated with the double-pseudoknot structure of the ribozyme though it lacks the P3 and P5 stem-loops. We additionally refer to the contacts T0 and T3 as tertiary contacts, where T0 consists of a *trans* Watson-Crick-Hoogsteen base pair U24 – A29 situated in loop L4 and T3 consists of base pair C15 – G19 situated at the truncated P3 step-loop (34), since we find them important for the ribozyme to assume its tertiary structure (Figure 1b). A biophysical study on the same ribozyme, including single-molecule Fluorescence Resonance Energy Transfer (smFRET) experiments, reported folding kinetics and self-cleavage activity in the presence of Mg^2+^ at various concentrations (36). Results from that study showed Mg^2+^ to facilitate the folding of the RNA although there was ambiguity around whether Mg^2+^ ions are involved in nucleolytic activity.

**Figure 1.**
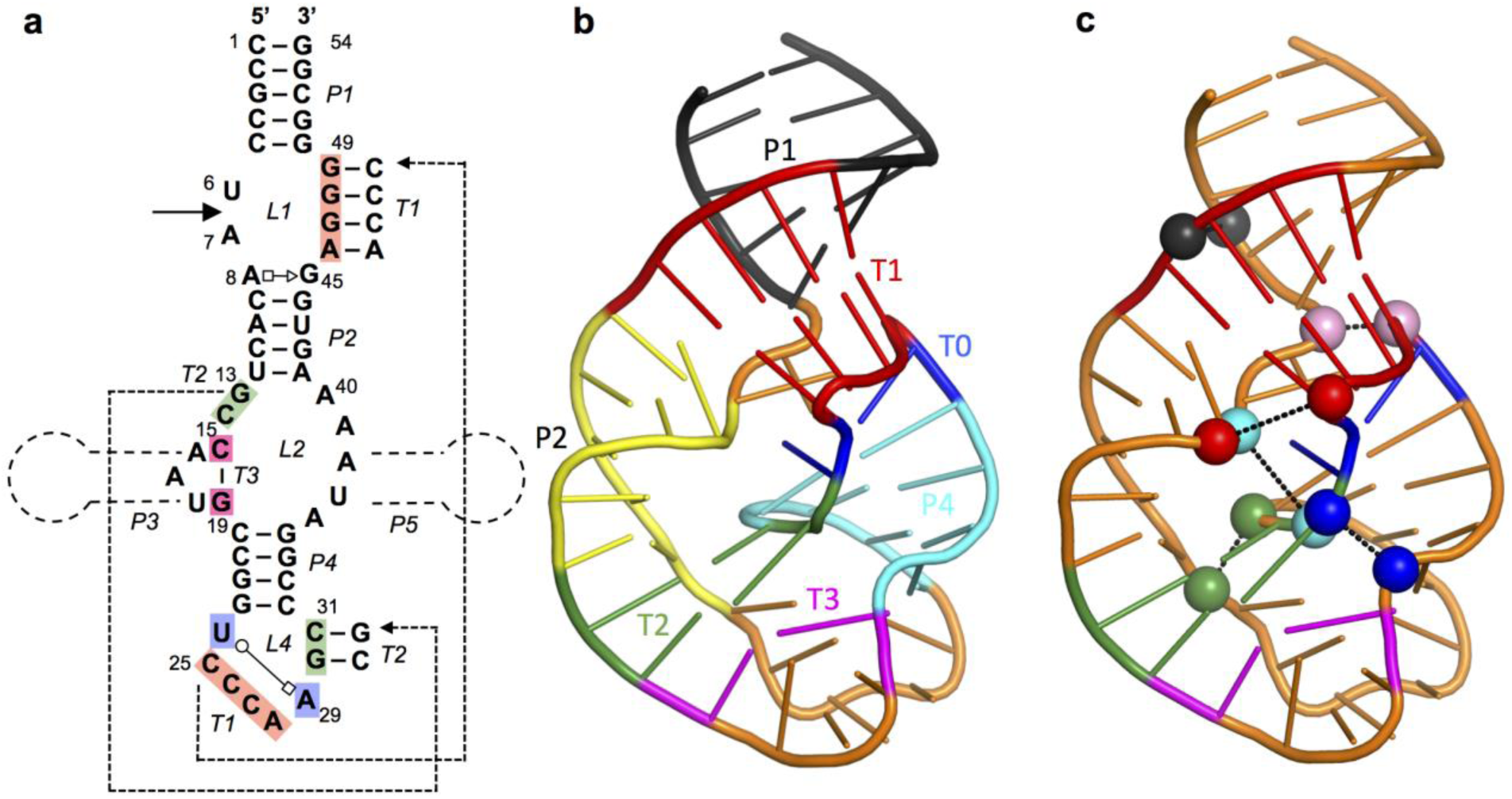
Structure of Twister ribozyme and spatially adjacent phosphate pairs. a) Twister Ribozyme secondary and tertiary contacts. The scissile bond is indicated with a solid arrow and the omitted P3 and P5 loops are indicated. b) Contacts are illustrated as cartoons colored as follows P1 – black, P2 – yellow, P4 – cyan, T0 – blue, T1 – red, T2 – green, T3 – magenta, rest of the RNA – orange. c) Twister ribozyme in its native state with spatially adjacent phosphate pairs that are present in the folded, tertiary structure (same-colored spheres connected with dashed line).

## Methods

### Potential of Mean Force Calculations

Potential of mean force (PMF) calculations along a folding pathway were performed using umbrella sampling in combination with the oscillating μ_ex_ GCMC-MD method. System preparation involved setting up four Twister simulation systems at 0, 10, 20 and 100 mM MgCl_2_ followed by subjecting each system to a classical MD simulation of 200 ns. The final coordinates from each of those simulations were used to generate conformations along the unfolding pathway. This was performed by rapidly unfolding the RNA based on the reaction coordinate going from 13 to 40 Å in 0.5 Å increments. Each of the subsequent 55 windows was then subjected to a 10 ns MD simulation in the presence of the umbrella potential at the respective RC distances. The final snapshots from these simulations were used to initiate the oscillating μ_ex_ GCMC-MD PMF calculation. This PMF calculation was performed in 1 Å increments from 13 to 40 Å yielding a total of 28 windows. Specific details for the different aspect of these calculations follow.

MD simulations of the Twister ribozyme were performed in a 90 □ cubic waterbox to accommodate the unfolded conformations at 4 different concentrations of Mg^2+^ (0, 10, 20 and 100 mM of MgCl_2_). The number of atoms in these systems is ∼68000. The 10 mM MgCl_2_ system corresponds to the number of Mg^2+^ ions in the Twister crystal structure (PDB 4OJI, 5 Mg^2+^ ions) and the initial positions of the ions were based on the crystal structure in this case. The two missing nucleotides in the crystal structure were added by using the internal coordinates in CHARMM (39). With 20 mM and 100 mM MgCl_2_ the Mg^2+^ were randomly placed in the simulation system. In addition to MgCl_2_, 100 mM KCl was used in all four systems. The systems were first minimized and equilibrated in CHARMM (39) with harmonic restraints on the backbone and base non-hydrogen atoms with force constants 1 and 0.1 kcal/mol/Å^2^, respectively, for 100 ps in the NPT ensemble with a timestep of 1 fs. CHARMM36 additive force field was used to model RNA and the ions (40-42). Water molecules were modeled by the CHARMM modified-TIP3P force field (43, 44). Smooth Particle Mesh Ewald method was applied for calculation of electrostatic interactions with a real-space cutoff of 12 Å (45). The Lennard-Jones potential was force switched to zero between 10 and 12 Å. A Monte Carlo anisotropic barostat was used to maintain pressure at 1 bar. The lengths of all covalent bonds that involve a hydrogen atom were constrained using the SHAKE algorithm (46). Equilibration was followed by production runs in the NPT ensemble using the Langevin integrator (1 bar, 30 °C) with no restraints for 200 ns in OpenMM (47, 48) with a time step of 2 fs.

Umbrella sampling MD was used to determine the potential of mean force (PMF) of breaking the tertiary contacts in Twister (49). The distance between the center of mass of two groups of nucleotides served as the reaction coordinate (Supporting Figure S1). The reaction coordinate was selected to globally lead to perturbation of the major tertiary contacts, T1 and T2, in Twister (Figure 1). The selection also allows for the relative positions of the fluorophores in the FRET study on Twister (36) to be monitored as a function of the RC. To generate unfolded conformations of the RNA the RC distance was gradually increased in 0.5 Å increments from 15 Å (initial average distance in the native conformation from the 200 ns MD simulations) to 40 Å (partially unfolded conformation), yielding a total of 51 windows. In addition, windows from 15 to 13 Å were generated and sampled in a similar fashion to assure that the true energy minima in the PMFs were identified. The windows were generated using CHARMM by running 10 ps MD simulations at each 0.5 Å step along the reaction coordinate with a force constant of 5000 kcal/mol/Å^2^ on the RC distance. During this stage the Watson-Crick (WC) base pairs forming the secondary interactions (P1, P2 & P4) were maintained by restraining the distances between the non-hydrogen atoms involved in all WC base pair hydrogen bonds. A force constant of 4 kcal/mol/Å^2^ was used to apply NOE restraints in CHARMM to maintain the distance between hydrogen bonding atoms of the bases within 2.7 Å to 3.0 Å. The final structure in each window was used as the starting structure for the next window. Each window was then simulated in the NPT ensemble for 10 ns in OpenMM (47) where the COM RC restraint was enforced using PLUMED using a force constant of 4.782 kcal/mol/Å^2^ (2000 kJ/mol/nm^2^) (50). No restraints were applied on the base pairs of secondary interactions and the RNA was allowed to propagate freely. The final frames from these 10 ns simulations were used to start the GCMC/MD simulation at alternate windows for the final PMF calculations. MD simulation conditions were the same in 200 ns production runs of the ribozyme in the native state.

### GCMC/MD protocol

A PME oscillating excess chemical potential (μ_ex_) GCMC algorithm was used to achieve enhanced sampling of the ions around the RNA (51) in conjunction with umbrella sampling to yield the final PMFs. At each window 2 separate PMFs were initially calculated. They each involved 5 cycles of oscillating μ_ex_ GCMC/MD with each cycle involving a resampling of the ion distribution in the simulation system using μ_ex_ GCMC, as described below, followed by 10 ns of NPT MD using the above protocol. From the 10 ns MD portions of the 10 oscillating μ_ex_ GCMC/MD cycles, the last 6ns of data from each of the 10 cycles were collected, yielding a total of 60 ns of sampling in each window from which the free energy profiles were calculated. Calculation of the PMF was performed using the Weighted Histogram Analysis Method (WHAM) and error analysis performed by block averaging over 5 12 ns blocks in each window and calculating the standard deviation. In total four systems (0, 10, 20 and 100 mM MgCl_2_) were subjected to umbrella sampling PMFs each involving 10 GCMC/MD cycles with 10 ns MD per cycle for each of the 26 windows yielding a total of 2.6 microseconds of MD for the Twister ribozyme.

The oscillating μ_ex_ GCMC involved a scheme to oscillate the μ_ex_ of the ions to overcome the low acceptance ratios and avoid using pre-hydrated Mg^2+^ ions as previously performed (52). In this protocol restricted Monte Carlo moves were used to resample the ion distribution by draining and refilling the system with Mg^2+^ ions, while K^+^ ions were added and drained simultaneously to neutralize the total charge on the system. For systems with no Mg^2+^ ions, the K^+^ and Cl^-^ ions were drained and refilled together to achieve redistribution of the K^+^ ions. One GCMC-MD cycle included seven stages: 1) Delete Mg^2+^ and insert K^+^ over 20000 MC steps; 2) Rotate and translate all ions and water for 80000 MC steps; 3) Repeat steps 1 and 2 until only one Mg^2+^ is left and the system is neutral; 4) Insert Mg^2+^ and delete K^+^ over 20000 MC steps; 5) Rotate and translate all ions and water for 80000 MC steps; 6) Repeat steps 4 and 5 until the Mg^2+^ concentration is reached and the system is neutral; and 7) minimize the entire system, including RNA, for 5000 Steepest Descent steps. Once resampling of the ion distributions was complete the system was then equilibrated for 100 ps of NVT MD followed by the 10 ns of NPT MD in OpenMM (47). In the PME/GCMC approach the μ_ex_ values to delete and insert the ions were adopted from the recent study of ionic hydration free energy and are shown in Supporting Table S2 (52).

### GFE maps

Occupancies of Mg^2+^ ions were calculated as the number of times a voxel (1 Å cubic unit of volume) around Twister ribozyme was occupied by Mg^2+^ during the simulation. At each window we analyzed 10000 frames and all frames were first aligned with respect to the backbone of the ribozyme. These occupancies were converted to “grid free energies” (GFE) according to G = −k_B_T ln(P), where P is the probability of occupying a voxel (1 Å cubic unit of volume) on the RNA surface relative to the voxel occupancy of the same species in bulk solution, k_B_ is the Boltzmann constant, and T is the temperature (303.15 K). More detailed description of this procedure is provided by Lemkul et al (32) and elsewhere (53).

## Results and Discussion

In the present study an enhanced sampling method for ion distributions is applied in combination with umbrella sampling to study atomic-level interactions of Mg^2+^ ions with fully and partially folded states of the Twister ribozyme. The protocol implemented includes GCMC simulations to redistribute the Mg^2+^, K^+^ and Cl^-^ ions in the simulation system. The GCMC is followed by minimization and MD simulations to allow the RNA, ions and water to relax, and to obtain conformational sampling of the RNA. The GCMC/MD approach overcomes the shortcomings of classical MD simulations due to the low exchange rates of Mg^2+^ ions with water and phosphate groups. Notably, the method achieves exchanges of ions around the phosphate groups at the inner-shell level, which is not currently feasible with classical MD. Validation of the method is detailed in Supporting Text 1. The combination of this approach in conjunction with umbrella sampling thus allows a direct relation of the atomic details of Mg^2+^-RNA interactions with the free energies associated with the stabilization of the RNA to be obtained as detailed in the remainder of this manuscript.

### Potentials of Mean Force as a Function of MgCl_2_ concentration

Twister ribozyme assumes a unique structure with a combination of secondary and tertiary interactions forming a double pseudoknot (34). The major tertiary contacts, T1 and T2, are long-range interactions that involve Watson-Crick (WC) base pairing. The nucleotides comprising the individual strands contributing to the two contacts are close in the primary sequence. Accordingly, the center of mass (COM) distance between two groups of nucleotides, regions 1 and 2, that approximately each represent half of the folded structure and each include the single strand regions that include T1 and T2 was chosen as the reaction coordinate (RC)(Supporting Figure S1). Regions 1 and 2 include the nucleotides U24 and G54, respectively, to which the individual fluorophores in the smFRET experiments were covalently linked (36). Along this RC a folding potential of mean force (PMF) of the Twister ribozyme was calculated using umbrella sampling with four different concentrations of Mg^2+^. To obtain the impact of Mg^2+^ on the PMFs, the umbrella sampling MD simulations at each RC were periodically stopped and the ion distribution in the system resampled using the GCMC protocol followed by equilibration and additional production umbrella sampling from which the full PMF was obtained. It is noted that this RC does not allow for sampling of the full range of unfolded states, as discussed below, but represents a computationally accessible 1-D RC allowing the sampling for unfolded states of the RNA to investigate their interactions with Mg^2+^.

The resulting PMFs for Twister at the four MgCl_2_ concentrations are shown in Figure 2a. The free energy surfaces show a sharp minimum at 15.5 Å, corresponding to the fully folded state. The free energy then increases rapidly to an inflection point at ∼20 Å, following which the free energy gradually rises out to the fully extended state at RC = 40 Å. From the minima out to 19 Å the surfaces at the four MgCl_2_ concentrations are similar. Beyond 19 Å the difference between the 0 and 100 mM MgCl_2_ surfaces are significant with the inflection point occurring at ∼23 kcal/mol for 100 mM MgCl_2_ and ∼30 kcal/mol for 0 MgCl_2_, with that difference being ∼7 kcal/mol throughout the 20 to 40 Å region of the PMF. The 10 and 20 mM MgCl_2_ system PMFs are between the 0 and 100 mM MgCl_2_ results beyond 20 Å out to 40 Å. To better understand these results with respect to experimental studies based on FRET and RNase T1 footprinting (36), analysis of the PMFs was undertaken specifically targeting the molecular phenomena directly related to those experiments.

**Figure 2.**
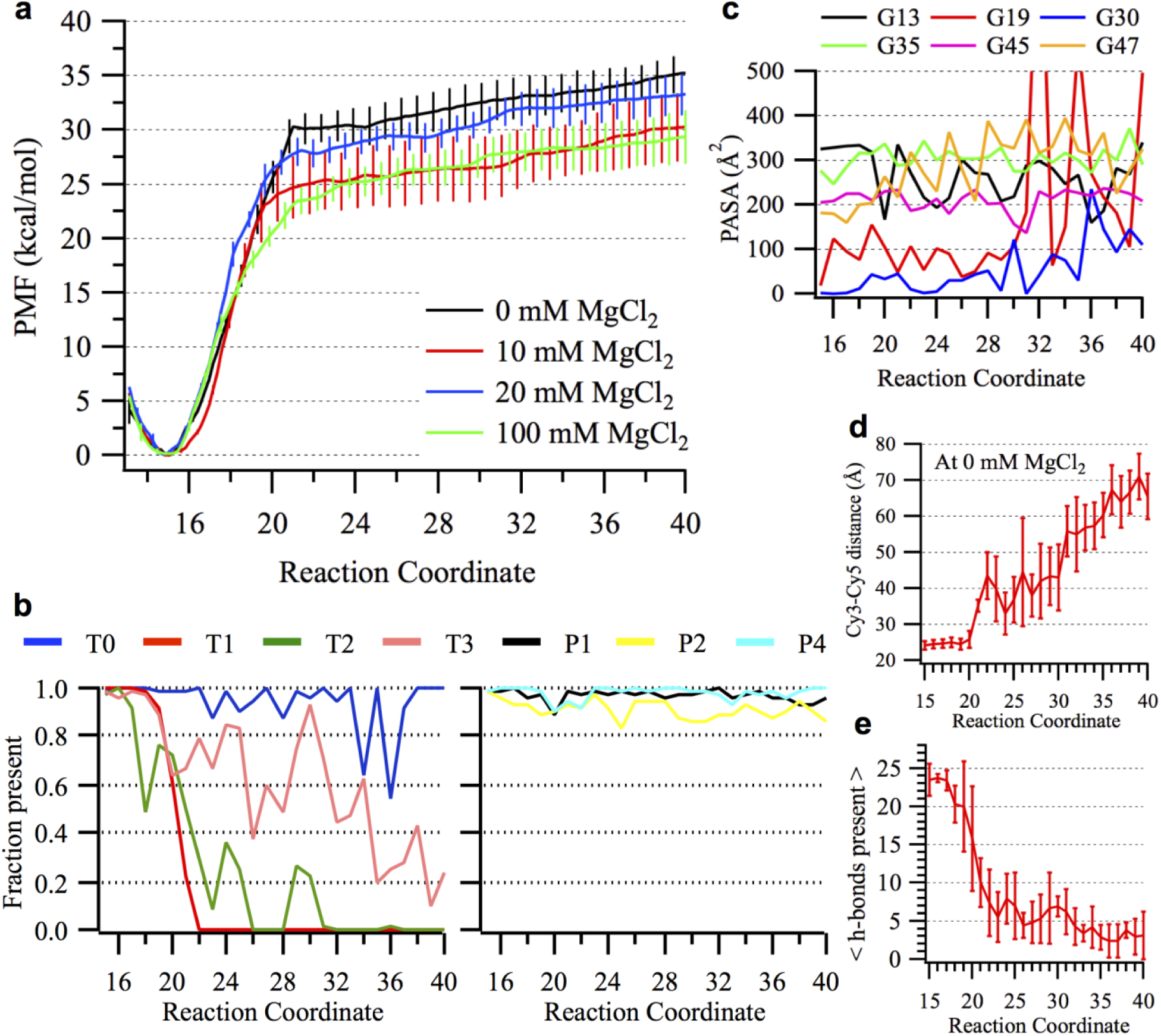
Analysis of Twister unfolding simulations. a) Potential of mean force (PMF) of Twister ribozyme at different concentrations of MgCl_2_ based on the GCMC/MD simulations. The reaction coordinate (RC) corresponds to center of mass distance between two groups of residues that contribute to the T1 and T2 tertiary interactions (Supporting Figure S1). Native state corresponds to RC = 15.5 Å. PMFs were calculated on RC windows at 1 Å intervals from 13 to 40 Å with 60 ns of sampling per window with error bars based on standard deviations from five 12 ns samples extracted from the full 60 ns of sampling (see Methods). b) The fraction sampled of tertiary (T0, T1, T2 and T3) and secondary (P1, P2, P4) contacts in Twister present at a given RC. Data represents averages over the four MgCl_2_ concentrations, with individual plots provided in Supporting Figures S5 and S6. A contact is present if the distance between the nucleotides is within 3 standard deviations of mean distance for the same contact calculated from the MD simulations in the native state. Fraction is calculated based on the number of frames a contact is present out of all frames analyzed. c) Protein accessible surface area (PASA) for the guanine nucleotides protected from cleavage by RNase T1 calculated at each window along the reaction coordinate with 0 mM MgCl_2_.

PASA is calculated as the solvent accessible surface area using a probe radius of 10 Å as previously described (54). d) Interfluorophore distance (IFD) based on distance between C4’ atoms of Ura 24 and Gua 54 along the reaction coordinate for Twister with 0 mM MgCl_2_. e) Number of Watson-Crick hydrogen bonds present between nucleotides involved in tertiary interactions along the reaction coordinate averaged over the four MgCl_2_ concentrations. Error bars indicate the standard deviation. A hydrogen bond was defined by donor-acceptor distance cutoff of 3.0 Å and donor-hydrogen-acceptor angle cutoff of 120°.

FRET experiments were based on the fluorophores Cy3 and Cy5; Cy3 was inserted between nucleotides U24 and C25 while Cy5 was connected to the 3’ termini via a ssDNA spacer (36). In the folded state the fluorophores are relatively close allowing for fluorescent energy transfer while upon unfolding the distance between the fluorophores increases leading to a loss of energy transfer. To model the spatial relationship of the fluorophores, distances between the C4’ atoms of Ura 24 and Gua 54 were measured from which the average and standard deviations were obtained for each window in the PMFs. Shown in Figure 2d are the average values of the “Interfluorophore distance (IFD)” along the RC for 0 MgCl_2_. Similar results were obtained with the other MgCl_2_ concentrations (Supporting Figure S2). Over RC values out to 20 Å and beyond the IFD does not change, with IFDs beyond 24 Å not occurring until RC = 22 Å at which point the PMFs have passed their inflection points, attaining energies of 23 kcal/mol or more. Accordingly, significant changes in the actual distance between the fluorophores that would lead to a loss of fluorescence will not occur until this distance along the PMF. In addition, given the length of the linkers on the fluorophores (Supporting Figure S2), including the region of ssDNA linking Cy5 to the RNA, significant unfolding of the RNA may be required to observe the change of fluorescence. This suggests that the FRET experiments may be reporting events that are occurring beyond the global minimum energy wells in the PMFs. Accordingly, interpretation of the present simulation results in the context of the FRET data will focus on the regions of the PMF beyond the inflection point at RC = 20 Å. In this region the 100 mM MgCl_2_ system is ∼7 kcal/mol more favorable than the 0 mM system with the 10 and 20 mM results intermediate to those values. This indicates that the presence of 100 mM MgCl_2_ lowers the free energy of Twister in these partially unfolded states. Supporting this interpretation is the presence of a small population of folded state at micromolar Mg^2+^ and a small population of unfolded state at and above 20 mM Mg^2+^ in the FRET experiments(36). The presence of small populations of the alternate states is consistent with a free energy difference of ∼ 7 kcal/mol versus a difference of over 20 kcal/mol between the global minima and the inflection at ∼20 Å in the PMF.

A similar interpretation of the experimental results on unfolding based on RNase T1 footprinting experiments may be made (36). Shown in Figure 2c are protein accessible surface areas over the guanine bases in Twister known to undergo hydrolysis. The individual plots with error analysis for each nucleotide are shown in Supporting Figure S3. While the accessibilities of different nucleotides vary, the values for the individual nucleotides do not increase significantly to well beyond the inflection point at 20 Å in the PMF. This indicates that increased hydrolysis being observed in the footprinting experiments is also not monitoring events associated with transitions from the global free energy minimum to the partially unfolded states (R = 20-40 Å), but rather associated with events beyond that region that correspond with larger unfolding events.

Consistent with the above model are β parameter results that are indicative of the region along the folding path where the transition state occurs leading to the fully unfolded state. With Twister β _fold_ = 0.37 and β _unfold_ = 0.72 indicating that the transition to the fully unfolded state observed in the FRET and footprinting experiments occurs closer to the unfolded state (36). In addition, the folding rate is under 0.1 s^-1^, which is not consistent with the over 20 kcal/mol minimum observed in the PMFs in Figure 2a (55). These results suggest that the experimental results are monitoring folding events that are occurring in the region beyond the inflection point at ∼ 20 Å in the PMF. In this scenario the increased population of the folded state observed in the FRET experiments in the presence of Mg^2+^ is consistent with the free energy of the 100 mM MgCl_2_ system being ∼7 kcal/mol more favorable that 0 mM MgCl_2_. However, beyond the inflection region in the PMF there is no indication of a transition to a further unfolded state, suggesting that the present RC is not sampling the full folding surface of the Twister RNA. Analysis of example conformations from the 100 mM MgCl_2_ PMF in Figure S4 of the supporting information show even at the longest RC = 40 Å, the RNA still contains a significant amount of Watson-Crick base pairing along with helical structure associated with the P1 loop. Loss of all or some of such interactions may lead to the unfolded states observed experimentally.

While the present PMF may not be representative of the full folding surface of Twister, the deep global minima may explain the ability of Twister to self cleave at very low Mg^2+^ concentrations. The presence of the deep minima, which are relatively insensitive to ion concentration, suggests that sampling of the folded state required for catalysis may occur in all conditions, which may be due to the background of 100 mM KCl (56) included in the present study to be consistent with the experimental study. Overall, the present results indicate a model where the global folded state observed experimentally is accessible even at low Mg^2+^ concentrations and that this conformation is related to the catalytic activity. However, the experimental FRET and RNase T1 footprinting studies are monitoring unfolding events at RC distances beyond the inflection point at 20 Å in the PMFs, with the present RC used to define the folding landscape not accessing the fully unfolded conformations. Additional studies are needed to address this issue.

Analysis of the tertiary and secondary contacts (Figure 1) averaged over all the four systems as a function of the RC is shown in Figure 2b, with results for the individual systems shown in Supporting Figure S5. In the global minima there is some loss of T2 tertiary interactions, the second largest tertiary contact, in the vicinity of RC = 17 to 18 Å as the free energy rapidly increases. The larger T1 interaction starts to be perturbed at RC 19 Å but it is not until beyond the inflection point in the PMF that the T1 interaction is lost, with the remaining T0 and T3 interactions largely maintained through the inflection point out to larger RC values. The secondary contacts are largely maintained throughout the PMFs (Supporting Figure S6), consistent with a report that secondary structures of RNA are independently stable of the tertiary interactions (57). In the P2 secondary interaction a cis-Hoogsteen-sugar edge pair between A8-G45 is lost during initial stages when T1 is being broken. In later stages with T1 fully broken, P2 is slightly compromised in order to stabilize the newly freed bases A46:G49. The inclusion of 100 mM background KCl in the present study likely contributes to the stabilization of the secondary interactions throughout the PMF.

To understand the contributions to the energy increase ranging from 23 to 30 kcal/mol upon moving from the global minima to the inflection point in the PMF the Watson Crick (WC) hydrogen bonds associated with the tertiary interactions were monitored. The tertiary interactions, T1, T2 and T3 correspond to 11, 6 and 3 hydrogen bonds, respectively, associated with 7 WCs pairs. Upon going from the fully folded state to the inflection point in the PMF approximately 80% of the T1 and T2 interactions are lost by 22 Å corresponding to a loss of 17 hydrogen bonds (Figure 2e). Considering that each WC hydrogen bond corresponds to a stabilization free energy ranging from -0.7 to -2 kcal/mol (58, 59) their loss corresponds to -12 to -34 kcal/mol indicating that the deep free energy wells in the PMFs are associated with perturbation of the hydrogens bonds associated with the T1 and T2 tertiary interactions. Analysis of traces of the phosphodiester backbone of Twister at various stages along the 100 mM MgCl_2_ PMF shows that at RC = 20 and 25 Å the overall 3D conformation of the RNA is maintained (Supporting Figures S4). The relatively small change in the overall conformation of the RNA upon going from the global minimum to the region of the inflection point in the PMF further supports the conclusion that the experimental FRET and RNase folding studies are not monitoring the initial loss of tertiary structure, but rather a larger scale unfolding event that is not being observed in the present calculations. This yields a scenario where the initial events in unfolding of Twister are dominated by local perturbations of hydrogen bond interactions participating in WC interactions present in the tertiary contacts while the overall structure of the ribozyme is maintained. Once these hydrogen bonds are broken, associated with the change in free energy of 23 to 30 kcal/mol seen in the PMFs, then larger scale unfolding of the RNA can occur associated with the gradual increase in free energy beyond the inflection point at 20 Å in the PMFs.

### Mg^2+^ distribution and non-sequential phosphate-phosphate interactions

While the present PMF doesn’t sample fully unfolded states, the energetic differences between the 0 and 100 mM MgCl_2_ concentrations beyond the PMF inflection point indicate that experimentally relevant partially unfolded states are being sampled. Accordingly, additional analysis focused on understanding the atomic details of the interaction of Mg^2+^ with Twister, including in both the folded state and partially unfolded conformations and how those interactions are leading to stabilization of the RNA. This analysis is based on the ability of the GCMC-MD protocol to sample a representative ensemble of Mg^2+^ ions around the RNA in both the folded and unfolded states.

Analysis of the distribution of Mg^2+^ ions around the RNA was initially undertaken for the folded state based on grid free energies (GFE) that are obtained directly from the ion probability distributions as described in the methods and previously performed (32). Figure 3 presents the distribution of Mg^2+^ around Twister from the RC = 15 Å PMF windows in the presence of 10, 20 and 100 mM MgCl_2_ along with the crystallographic ion positions. It is evident that the simulations produce distributions that recapitulate the crystallographic locations of Mg^2+^ even at the lower MgCl_2_ concentration. It shows that with decreasing concentration the Mg^2+^ ions still sample sites observed in the crystal structure along with additional regions. Figure 3c shows that even at 10 mM MgCl_2_, Mg^2+^ ions are specifically distributed at crucial intersections, the majority of which correspond to the specific phosphate pairs discussed below. However, at the lower MgCl_2_ concentrations it is likely the convergence of the ion distribution may require additional sampling, which may impact the free energy surfaces shown in Figure 2a. Additional tests are required to address this issue.

**Figure 3.**
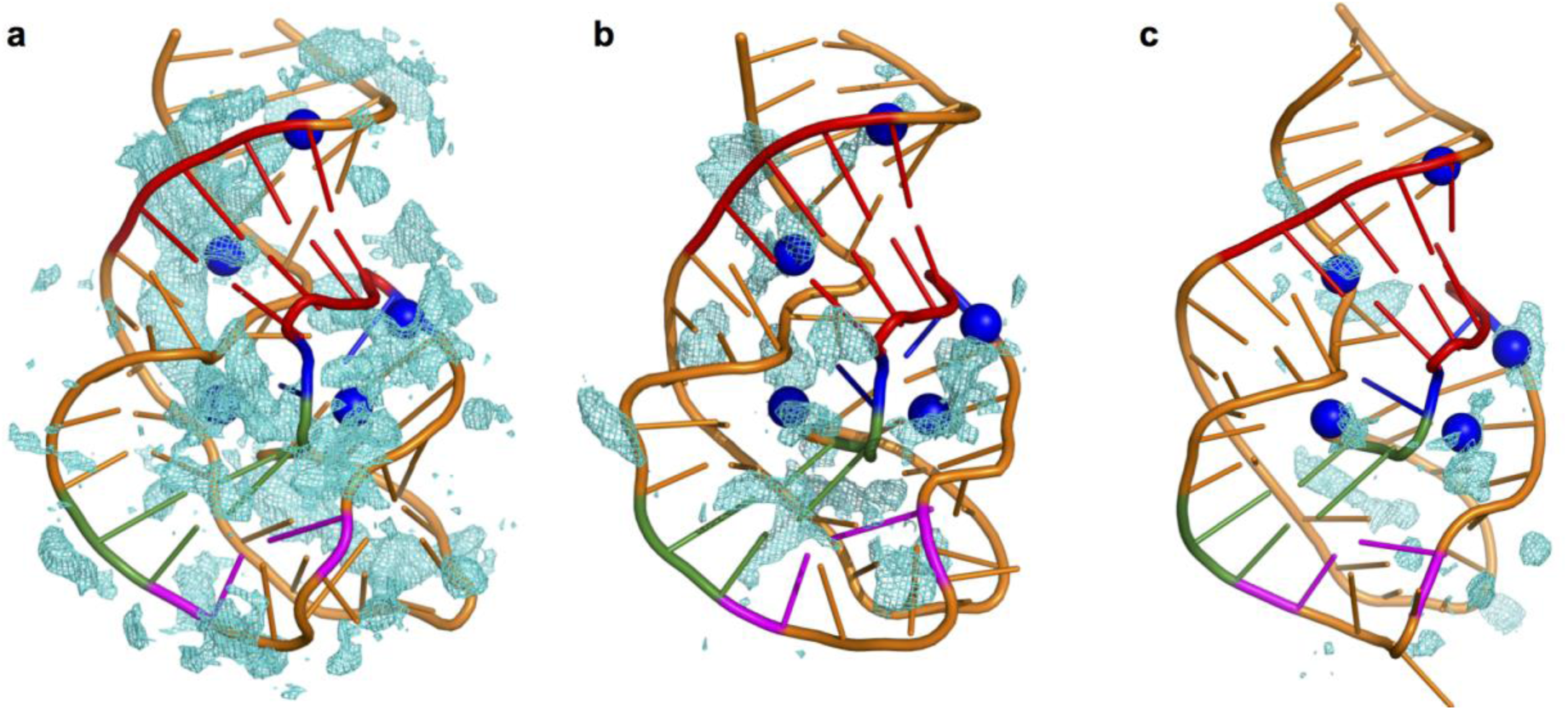
Occupancy maps for Mg^2+^ ion distribution around native conformation of the Twister ribozyme from the GCMC-MD simulations. a) 100 mM MgCl_2_ b) 20 mM MgCl_2_ and c) 10 mM MgCl_2_. The cutoff for GFE is -2 kcal/mol. The crystallographic positions of Mg^2+^ binding sites are shown in blue spheres after aligning the structures.

Further analysis of interactions of the RNA with Mg^2+^ along the reaction coordinate focused on the presence of spatially adjacent non-bridging phosphate oxygens (NBPOs) of non-sequential nucleotides (*i.e*. phosphate moieties separated by one or more phosphates in the primary sequence) and their spatial relationship to Mg^2+^. Phosphate-Mg^2+^ interactions, including both direct and indirect coordination, may stabilize both the folded and partially unfolded states by facilitating short phosphate-phosphate interactions. Non-sequential NBPO pairs with a high probability of being within 9 Å of each other were identified as pairs that may simultaneously interact with Mg^2+^ through outer shell interactions (Figure 4). The cutoff of 9 Å is defined based on radial distribution functions (RDF) of the distance between NBPO and Mg^2+^ ions (Supporting Figure S7) that show peaks at 1.95 Å, 4.2 Å and 6 Å corresponding to direct-interaction (Inner-Shell-Dehydrated: ISD), indirect-interaction (Outer-Shell-Dehydrated: OSD) and diffuse-interaction (Non-Dehydrated: ND), respectively (8). Here, we focus on the ISD and OSD interactions, which have been classified as strong interactions.

**Figure 4.**
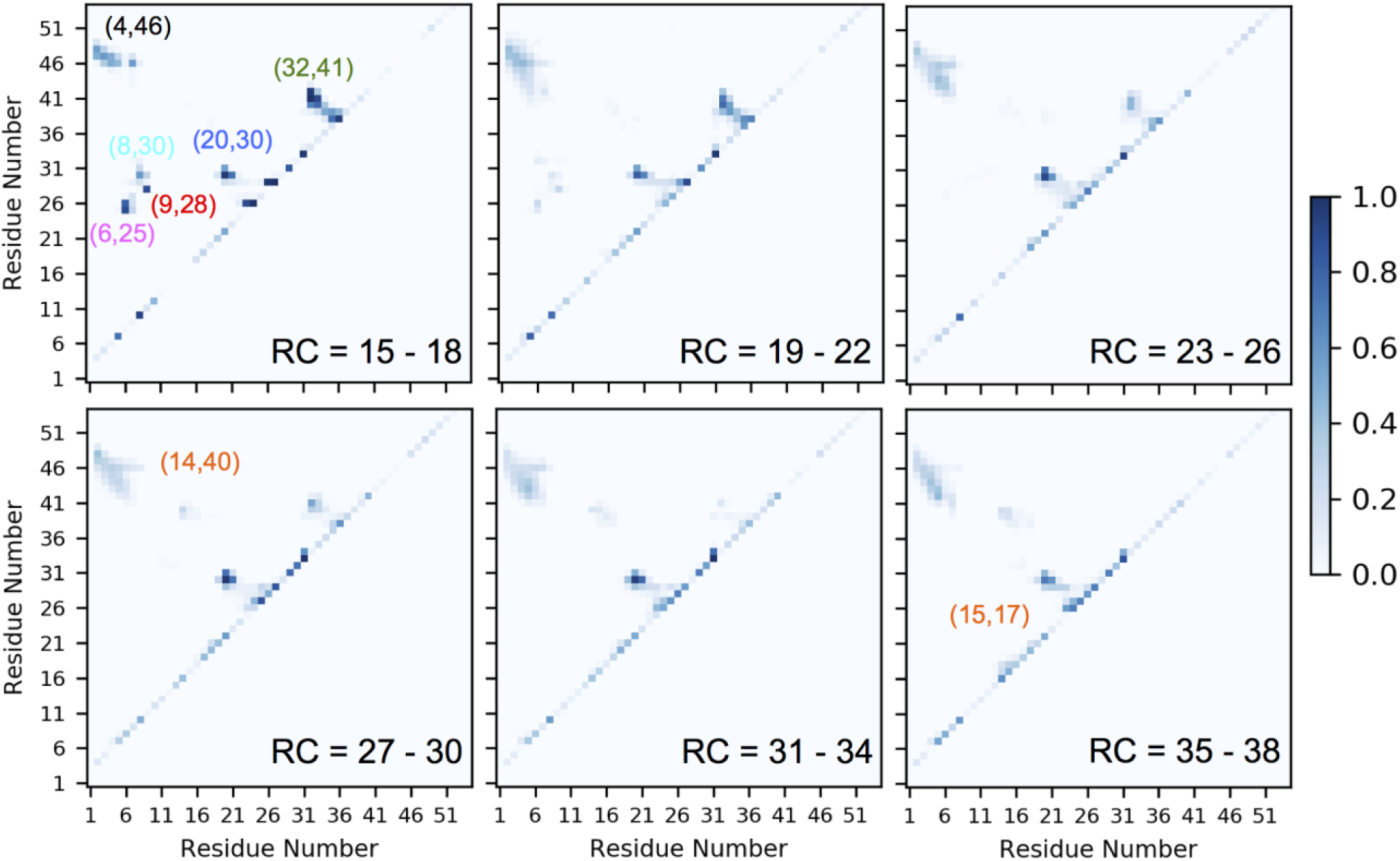
NBPO probability matrices from NBPO distance analysis for Twister from the GCMC-MD PMF at 100 mM MgCl_2_. The probabilities were averaged over groups of 4 RC windows. RC = 15 – 18 corresponds to the fully folded structure. The interactions discussed in the text have been specified with circles.

To identify interacting non-sequential NBPOs, adjacent NBPO probability analysis was performed for selected portions of the PMF by merging the data for four consecutive windows (Figure 4). Analysis of Figure 4 shows the presence of non-sequential NBPO pairs that are adjacent to each other that are far apart in the primary sequence as well as a number of NBPO pairs that are adjacent to the diagonal associated with phosphates separated by one or more nucleotides. Table 1 defines six regions of non-adjacent NBPO interactions that occur in the folded state that are identified in the top-left panel of Figure 4. These are shown in Figure 1c using the same color scheme as used for the labels in Table 1. Of these a number are situated around the T1 and T2 tertiary interactions (Table 1). Notable of the pairs present in the folded state is the pair C9-A28 (Figure 1c, red spheres) in which A28 is situated at the end of T1, such that this pair coming together facilitates base-pair formation in T1. These NBPOs are exposed to the solvent allowing for Mg^2+^ to simultaneously coordinate with both of them. Additionally, the U6-C25 pair (Figure 1c, pink spheres) is situated on the other side of T1 further indicating the importance of bringing phosphates together to stabilize the pseudoknot structure of the ribozyme. Between G3:C5 and A46:G47 (Figure 1c, C4-A46 black spheres) there is a range of sites where Mg^2+^ ions may interact with more than 2 NBPOs. Formation of these pairs appear to stabilize the helical twist of P1 secondary motif while stabilizing nucleotides involved in T1 that may facilitate formation of that contact. The pairs A8-G30 and A8-C31 (Figure 1c, cyan spheres) provide an encapsulated space for Mg^2+^ that bridges nucleotides on T1 and T2. NBPO pairs between C32:C33 and A41:G42 (Figure 1c, C32-A41 green spheres) are adjacent to and appear to stabilize the T2 interaction. Finally, the C20-G30 pair (Figure 1c, blue spheres) is within 9 Å throughout the majority of the PMF and appears to stabilize nucleotides in the hairpin loop on the end of P4 that participate in both T1 and T2 as well as stabilizing the T0 (U24-A29) and T3 (C15-G19) interactions.

**Table 1.** List of pairs that show high probabilities of their NBPOs being within 9.0 Å in the folded state.

|  | Pairs with NBPO correlation | Adjacent tertiary interaction |
| --- | --- | --- |
| ●● | C4 – A46 | T1 |
| ●● | C9 – A28 | T0, T1 |
| ●● | A8 – G30 | T1 |
| ●● | U6 – C25 | T0, T1 |
| ●● | C32 – A41 | T2 |
| ●● | C20 – G30 | T2, T3 |

Figure 5 shows the distribution of Mg^2+^ ions around the phosphate pairs listed in Table 1 using GFE maps at RC = 15 Å in the 100 mM MgCl_2_ system. The presence of highly favorable regions of Mg^2+^ sampling directly adjacent to or between all the NBPO pairs is evident. For example, a large region is sampled around the pair 4-46 where phosphates are lined in parallel fashion with NBPOs facing each other. In general, the presence of Mg^2+^ in the regions surrounding the interacting NBPO pairs will stabilize these interactions, which will lower the free energy of the partially unfolded states. In addition to the NBPO pairs discussed above, there are high probability pairs adjacent to the diagonal of the matrix in top left panel of Figure 4. These include 5-7, 8-10, 24-26, 27-29, 29-31, 31-33, and 36-38. These represent kinks in the backbone of the ribozyme that may play an important role in the formation of tertiary structure. It has been reported that such kinked structures are stabilized by divalent metal ions (60).

**Figure 5.**
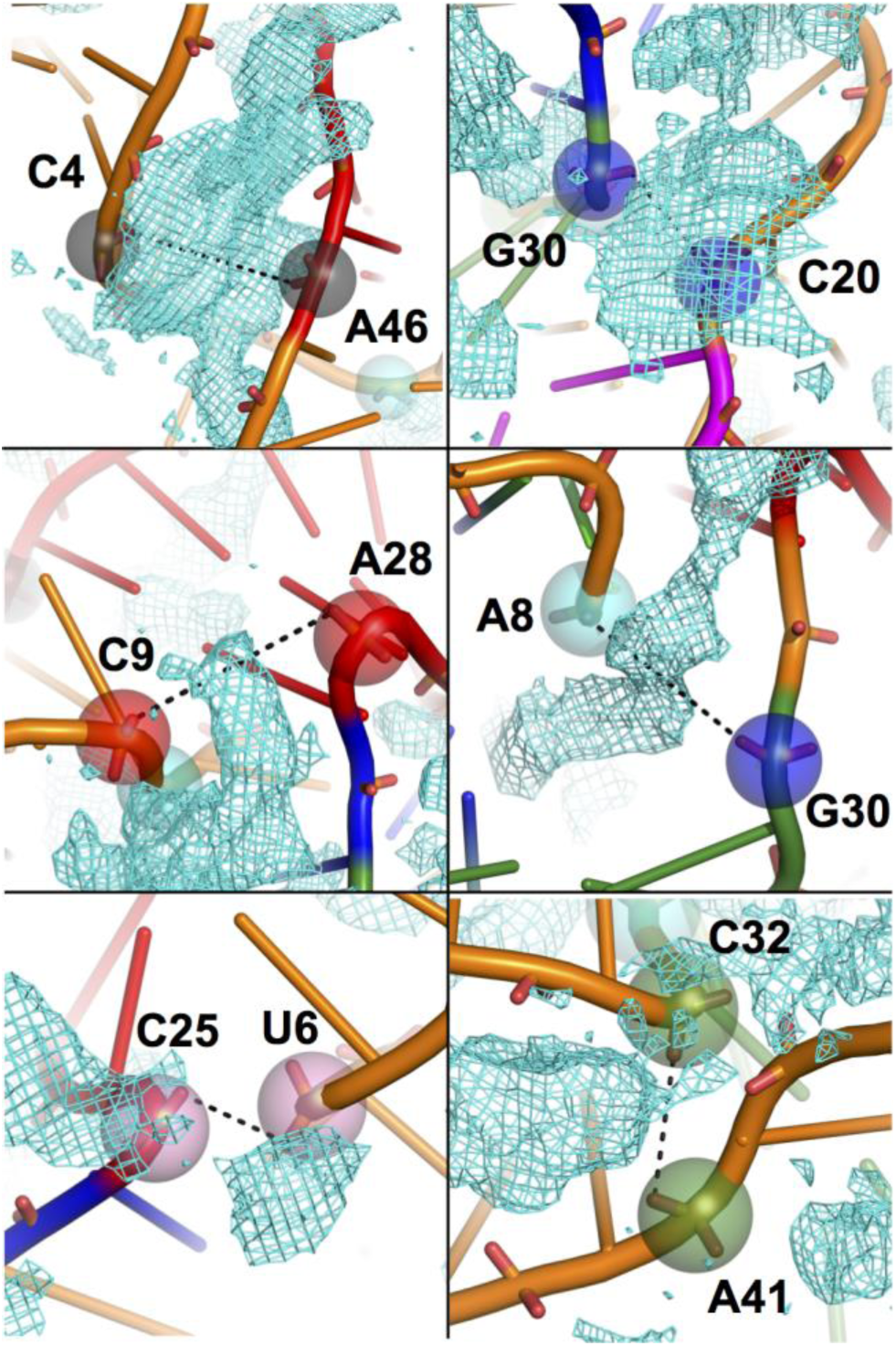
Distribution of Mg^2+^ ions around the NBPOs of residue pairs. From the RC = 15 Å native state of Twister ribozyme (cartoon) from the 100 mM MgCl_2_ PMF. The phosphates are represented as same-colored transparent spheres. GFE maps are illustrated as cyan mesh at -2 kcal/mol cutoff.

At higher RC values a number of the pairs present in native state are lost while two new pairs appear. Pairs that are lost correspond to pairs whose individual partners are assigned to the two groups used to define the RC. For example, the 6-25, 8-30, and 9-28 NBPO pairs are lost. New interacting pairs at long RC values include the 15-17 pair that occurs at RC values greater than 30 Å and the 14-40 pair that first occurs in the 27-30 RC portion of the PMF.

The NBPO probability analysis was also performed for Twister with 0 mM MgCl_2_ (Supporting Figure S8). In the absence of Mg^2+^ ions, near the folded state (RC = 15 to 18) the pattern of NBPO pair is similar to that at 100 mM MgCl_2_, with the 9-28, 8-30, 6-25 and 32-41 pairs being present. However, some differences occur at larger RC values. The group of NBPOs near the 4-46 pair shows lower probabilities with the interaction pattern shifted and the interaction between 15-17 pair at RC = 35-38 is not observed. The similarity of the NBPO pairs in the folded state is consistent with experimental data indicating the ability of Twister to fold and self cleave at concentrations approaching 0 Mg^2+^ as well as with the present PMF (Figure 2a), while the difference at the larger RC values suggest a role of the ion in stabilizing the partially unfolded states.

To identify Mg^2+^ ions in the vicinity of the NBPO pairs along the PMF the probability of Mg^2+^ being present within 6.5 Å of the NBPO atoms of both phosphates in each pair along the RC was calculated. Data are presented for the pairs in Table 1 along with the 14-40 and 15-17 pairs seen in the partially unfolded states. Shown in Figure 6 are those values as a function of the RC along with the fraction of the NBPO pair that is within the 9 Å cutoff. It is evident that whenever the NBPOs are close to each other there is a significant probability of Mg^2+^ ion being in contact with both of them. The collection of NBPO pairs designated by 4-46 (Figure 4) was present at fraction probabilities of 0.2 or more along the RC with similar probabilities of Mg^2+^ in their vicinity. While this interaction is present for the full range of the PMF, it occurs at higher probabilities in the vicinity of the folded state with the Mg^2+^ probability following that trend. With pairs 9-28, 8-30 and 6-25, which involve nucleotides participating in T1, Mg^2+^ ions are interacting with both members while those interactions are present out to RC = 22 Å. The pair 32-41, which is situated around tertiary contact T2, also has Mg^2+^ extensively coordinated with this NBPO pair. As discussed above, pair 20-30 is maintained along the full PMF with Mg^2+^ ions consistently interacting with the pair. Notably, a similar pattern is observed with the NBPO pairs not present in the folded state. The presence of both the 14-40 and 15-17 pairs in the unfolded states along the PMF is strongly correlated with interactions with Mg^2+^. Such transient interactions that are facilitated by Mg^2+^ may be important for guiding the RNA along the folding pathway, as previously discussed (18, 25), though further all-atom simulation studies are needed to directly address this possibility.

**Figure 6.**
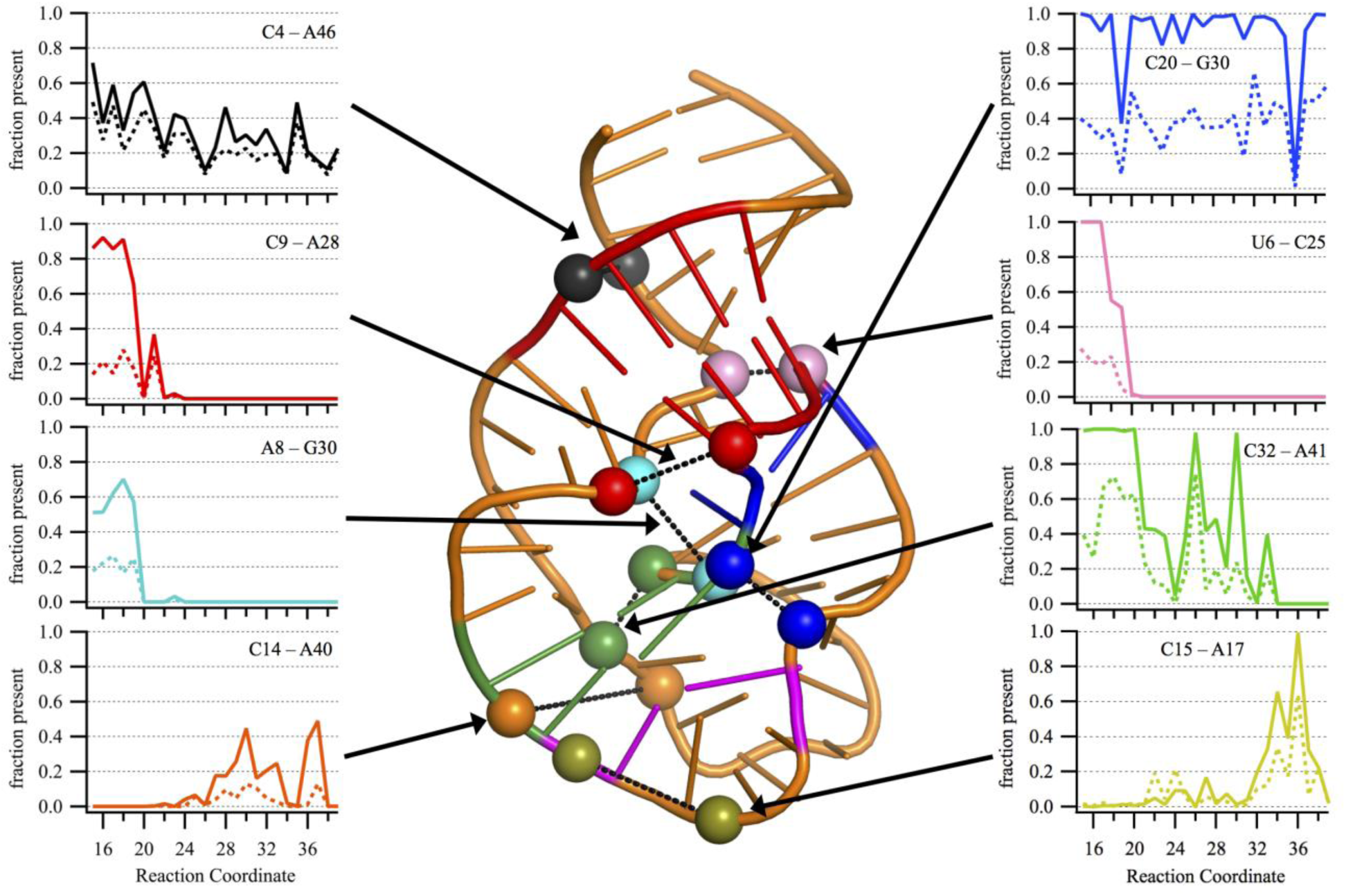
Adjacent NBPO pairs in contact with Mg^2+^ ions at 100 mM MgCl_2_. The probability of different pairs of nucleotides with NBPOs within 9 Å (Solid line) and the probability of a Mg^2+^ ion within 6.5 Å of NBPOs of both phosphates coordinating the two NBPOs (Dashed line) along the reaction coordinate. The arrows point out the positions of NBPO pairs in the Twister structure shown in cartoon and the phosphate atoms shown as spheres with similar colors. Plots for 20mM and 10 mM MgCl_2_ are provided in Supporting Figure S9.

Interesting trends are seen with NBPO pairs that impact the T0 and T3 tertiary contacts. T0 is present throughout the majority of the PMF, though occurs at a lower probability in the most unfolded states (Figures 2b). This contact, which occurs in the L4 loop, is associated with the 20-30 NBPO pair maintained throughout the PMF. A similar trend is present with the T3 tertiary interaction (15-19 bulge) for which sampling at low levels occurs in the most unfolded states sampled in this study and gradually increasing at smaller RC values (Figure 2b). Analysis of the NBPO probability matrices shows the presence of a 15-17 interaction near the diagonal at RC=35-38, and Figure 6 shows that Mg^2+^ ions are present around this pair. It suggests that this interaction, which is associated with a kink in the RNA, facilitates formation of the T3 contact.

To quantify the impact of the presence of Mg^2+^ on the NBPO pairs the overall probabilities of the presence of those pairs was calculated by integrating over all the individual pair probabilities excluding those in adjacent phosphates. The results from this analysis are presented in Table 2. Comparison of the results for 100 and 0 mM show the probabilities to be systematically higher throughout the PMF at 100 mM MgCl_2_, with 10 and 20 mM values typically falling between the 100 and 0 mM values consistent with the PMFs shown in Figure 2a. This difference indicates that while the majority of NBPO pairs are present in 0 MgCl_2_, the presence of Mg^2+^ has an overall stabilizing effect on those interactions. This is consistent with the model of Mg^2+^ stabilizing the repulsion associated with phosphate-phosphate interactions occurring in the tertiary structure (7, 61). Thus, the presence of Mg^2+^ neutralizing phosphate-phosphate repulsion associated with tertiary interactions leads to an overall increase in the sampling of the non-sequential NBPO pairs in Twister. This increased sampling is suggested to contribute to the lowering of the free energy of the RNA as seen in the differences in the PMFs beyond the inflection point where the 100 mM MgCl_2_ system is consistently 7 kcal/mol lower than the 0 mM MgCl_2_ system.

**Table 2.**
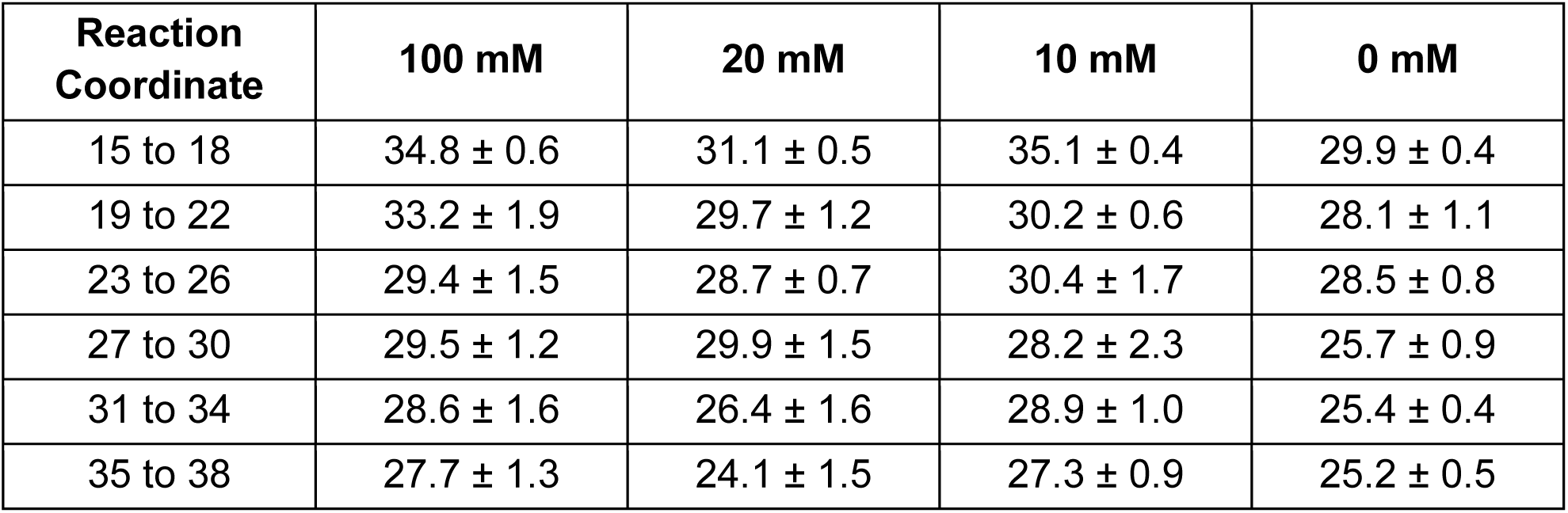
Summed probability of the non-sequential NBPO pairs being within the 9.0 Å cutoff at various stages of unfolding for systems at different Mg^2+^ concentrations. The analysis involves all phosphate-phosphate pairs that are not on adjacent sequential nucleotides in the RNA.

| Reaction Coordinate | 100 mM | 20 mM | 10 mM | 0 mM |
| --- | --- | --- | --- | --- |
| 15 to 18 | $34.8 \pm 0.6$ | $31.1 \pm 0.5$ | $35.1 \pm 0.4$ | $29.9 \pm 0.4$ |
| 19 to 22 | $33.2 \pm 1.9$ | $29.7 \pm 1.2$ | $30.2 \pm 0.6$ | $28.1 \pm 1.1$ |
| 23 to 26 | $29.4 \pm 1.5$ | $28.7 \pm 0.7$ | $30.4 \pm 1.7$ | $28.5 \pm 0.8$ |
| 27 to 30 | $29.5 \pm 1.2$ | $29.9 \pm 1.5$ | $28.2 \pm 2.3$ | $25.7 \pm 0.9$ |
| 31 to 34 | $28.6 \pm 1.6$ | $26.4 \pm 1.6$ | $28.9 \pm 1.0$ | $25.4 \pm 0.4$ |
| 35 to 38 | $27.7 \pm 1.3$ | $24.1 \pm 1.5$ | $27.3 \pm 0.9$ | $25.2 \pm 0.5$ |

To investigate the impact of Mg^2+^ on the sampling of the individual NBPO pairs the summed probabilities were obtained over the different regions of non-sequential NBPO pairs shown in Figure 4 at 100 mM vs. 0 mM MgCl_2_ and the differences determined. In addition, the contributions from all pairs adjacent to the diagonal were obtained. The differences between the two concentrations are shown in Table 3. As may be seen, in the vicinity of the folded state and the region leading to the inflection point in the PMF a number of pairs are more populated in 100 mM MgCl_2_ including 8-30, 32-41, 4-46 and 20-30. The largest difference occurs with the 4-46 interactions with that increase occurring throughout the PMF. More sampling is also seen with the 15-17 pair at the RC values beyond 30 Å. For the interactions adjacent to the diagonal the difference are relatively small, being both higher and lower depending on the region of the PMF. Interestingly, for some non-sequential NBPO pairs there is a decrease in the probability of sampling at 100 mM MgCl_2_ indicating that while Mg^2+^ leads to an overall increase in sampling of non-sequential NBPO pairs, there is variability in the specific contributions at an atomic level.

**Table 3.**
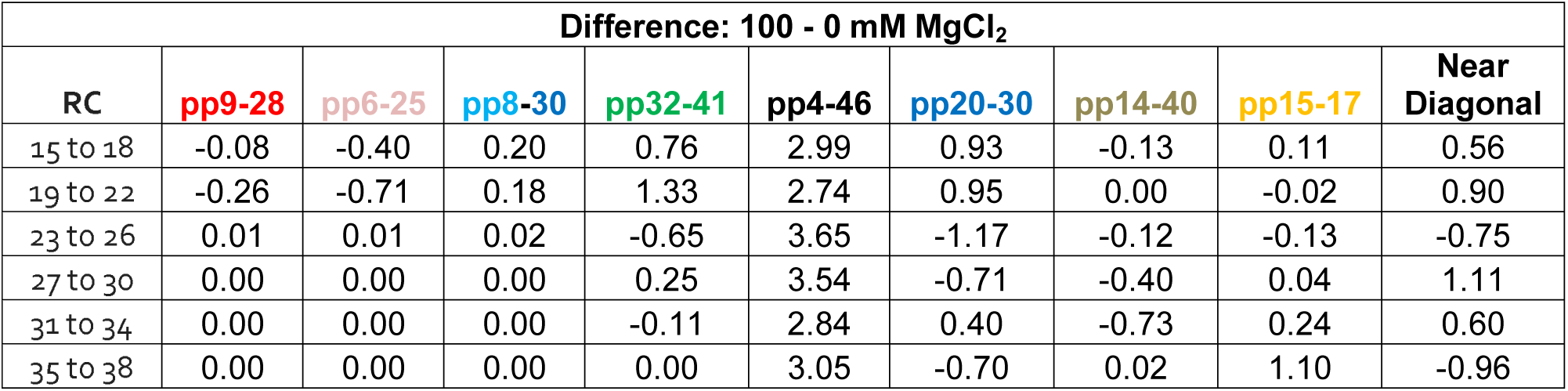
Difference between the summed probabilities at 100 and at 0 mM MgCl_2_ for the individual non-sequential NBPO pairs and for the non-sequential pairs adjacent to the diagonal on Figure 4. The near diagonal values were calculated based on the total values shown in Table 2 minus the sum of the NBPO pairs listed in Table 3. Absolute values are shown in Supporting Table S1a and standard deviations in the differences are shown in Supporting Table S1b.

| Difference: 100 - 0 mM $\text{MgCl}_2$ | | | | | | | | | |
| --- | --- | --- | --- | --- | --- | --- | --- | --- | --- |
| RC | pp9-28 | pp6-25 | pp8-30 | pp32-41 | pp4-46 | pp20-30 | pp14-40 | pp15-17 | Near Diagonal |
| 15 to 18 | -0.08 | -0.40 | 0.20 | 0.76 | 2.99 | 0.93 | -0.13 | 0.11 | 0.56 |
| 19 to 22 | -0.26 | -0.71 | 0.18 | 1.33 | 2.74 | 0.95 | 0.00 | -0.02 | 0.90 |
| 23 to 26 | 0.01 | 0.01 | 0.02 | -0.65 | 3.65 | -1.17 | -0.12 | -0.13 | -0.75 |
| 27 to 30 | 0.00 | 0.00 | 0.00 | 0.25 | 3.54 | -0.71 | -0.40 | 0.04 | 1.11 |
| 31 to 34 | 0.00 | 0.00 | 0.00 | -0.11 | 2.84 | 0.40 | -0.73 | 0.24 | 0.60 |
| 35 to 38 | 0.00 | 0.00 | 0.00 | 0.00 | 3.05 | -0.70 | 0.02 | 1.10 | -0.96 |

While Mg^2+^ overall stabilizes Twister as well as other RNAs, the results in Table 3 indicate a more complex picture. Several pairs, including 8-30, 32-41, and 20-30 are stabilized in the vicinity of the inflection point in the PMF as well as in the fully folded states as expected. The 4-46 interaction is significantly stabilized throughout the PMF, indicating its impact on the stabilization of the RNA by Mg^2+^ in the full range of conformations studied. In contrast, decreases in the sampling of adjacent NBPO pairs occurs with 9-28 and 6-25 in the inflection and fully folded regions of the PMF while decreases at longer RC values occur with the 32-41, 20-30 and 14-40 pairs. With 14-40 the decreased sampling of close interactions is consistent that that pair not being present in the folded state, such that its destabilization would facilitate folding. With the 32-41 and 20-30 pairs the destabilization in the RC=23 to 26 region may help to avoid formation of tertiary interaction too early in the folding process, thereby facilitating the overall folding process. The largest decrease in the sampling of the 9-28 and 6-25 pairs occurs around the inflection point on the PMF at RC=19 to 22 Å. As these pairs are directly adjacent to the T1 tertiary contact, their destabilization may facilitate proper formation of the WC hydrogen bonds that dominate that tertiary contact. Notably, Mg^2+^ is sampling in the vicinity of these pairs (Figure 5) indicating that despite the presence of Mg^2+^ the increased sampling of other NBPO pairs occurs at the expense of these interactions consistent with a push-pull mechanism for the stabilization of RNA by Mg^2+^.

To further investigate the relationship of the 9-28 and 6-25 NBPO pairs to the T1 tertiary contact, correlation analysis was undertaken between the individual NBPO pairs and the number of T1 WC-associated hydrogen bonds in the region of the inflection point in the PMF (RC = 19 to 21, Table S1c). The Pearson correlation coefficients in the case of the 9-28 NBPO pair were -0.35 and -0.62 in 0 and 100 mM MgCl_2_, respectively, while with the 6-26 NBPO pair the values were -0.37 to -0.65, respectively. Plots of the NBPO distances versus number of T1 WC hydrogen bonds are presented in Supporting Figure S10. These results show that the shorter NBPO distances correspond to a larger number of T1 hydrogen bonds, as expected given the spatial relationship of the two NBPO pairs to the T1 tertiary contact (Figure 1). Interestingly, the increase in the magnitude of these correlations in the presence of Mg^2+^ indicates how the ion influences the communication between these two classes of interactions. However, the decrease in the sampling of shorter NBPO distance in the presence of Mg^2+^ (Table 3) indicates that the formation of the T1 hydrogen bonds is actually decreased due to the presence of Mg^2+^. This further indicates a complex push-pull type of mechanism where the presence of Mg^2+^ overall favors the tertiary structure but the lowered sampling of short 9-28 and 6-25 NBPO pairs in the inflection region of the PMF where the tertiary contacts are initially forming may allow for the hydrogen bonds to sample a wider range of conformational space. Such additional sampling would allow for formation of the correct pattern of hydrogen bonds associated with the WC interactions that occur in the fully folded state.

## Conclusions

The central observation from the present calculations is the overall stabilization of the non-sequential NBPO interactions pairs by Mg^2+^, an observation made in course-grained simulations studies of the *Azoarcus* ribozyme (18, 25). Notably, in the present study it is shown that Mg^2+^ leads to increased probability of the sampling of specific non-sequential NBPO pairs offering direct evidence of Mg^2+^ contributing to the energetic stabilization of the RNA observed in the PMFs. The most notable such effect occurred in the vicinity of pair 4-46. The region is involved in stabilization of the P1 helix along with the nucleotides that participate in the T1 tertiary contact. This suggests the Mg^2+^ is specifically interacting with the RNA to stabilize the conformation of this region thereby facilitating formation of the T1 contact upon full folding of the RNA. Previous studies have reported conflicting results on the importance of this stem in folding and activity (34, 37, 38). Interestingly, Mg^2+^ is not seen in this region of Twister in the crystallographic structures (34, 38).

Other NBPO pairs are stabilized or destabilized to varying extents consistent with Mg^2+^ facilitating RNA folding and stability in a sequence specific fashion. Decreased sampling of specific non-sequential NBPO pairs in the presence of Mg^2+^ is a somewhat surprising result (Table 3). For example, in the vicinity of the inflection region of the PMF and in the folded states the sampling of the 9-28 and 6-25 pairs is decreased by Mg^2+^ as also occurs in the partially unfolded states with the 14-40 pair. This shows that the specific effects of Mg^2+^ include destabilization of selected non-sequential NBPO pairs as other phosphate-phosphate interactions are being stabilized in a push-pull mechanism. In the case of 14-40, as this interaction is not present in the folded states, its destabilization would facilitate access to those folded states. In the case of 9-28 and 6-25 the results indicate that partial destabilization of the pairs facilitates proper formation of the hydrogen bonds associated with the WC interactions of the T1 tertiary contact such that Mg^2+^ allows Twister to break and form the hydrogen bonds associated with the T1 tertiary contact. This appears to contribute to the smooth transition in the PMFs around inflection point in the presence of Mg^2+^ versus in its absence where a sharp transition occurs (Figure 2a). These results indicate a push-pull scenario of Mg^2+^ stabilization of RNA where, despite the general neutralization of phosphate-phosphate repulsion, the nature of the Mg^2+^ interactions leads to lowered sampling of specific non-sequential phosphate pairs that are being sacrificed to yield an overall stabilization of the RNA. While we anticipate that the identified push-pull mechanism contributes to the stabilization of other RNAs by Mg^2+^, given the unique features of Twister where Mg^2+^ concentrations approaching zero still allow for self cleavage, it would appear that the role of Mg^2+^ in RNA structure and function varies for different systems.

## Author Contributions

ADM Jr. conceived the overall project. AAK and ADM Jr. designed the methods and models for simulations. AAK built and performed the simulations. AAK and ADM Jr. analyzed the data and wrote the manuscript.

## Conflict of Interest

ADM Jr. is co-founder and CSO of SilcsBio LLC.

### Acknowledgements

We thank the National Institutes of Health [GM131710] for financial support for this work and the Computer-Aided Drug Design Center at the University of Maryland Baltimore for computing time. We thank Dr. Sarah Woodson and Dr. Darrin York for helpful discussions.

## Supporting Citations

References (62-64) appear in the Supporting Material.

