## Supplemental figures, tables and text for "Mg^2+^ Impacts the Twister Ribozyme through Push-Pull Stabilization of Non-Sequential Phosphate Pairs"

A. A. Kognole and A. D. MacKerell, Jr.

### SUPPORTING FIGURES

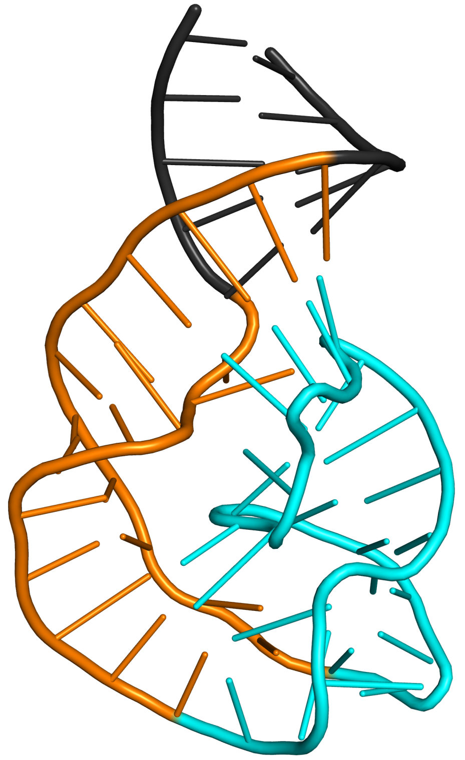

Figure S1. Reaction coordinate was calculated as the distance between the centers of mass of the non-hydrogen atoms in group1 (orange cartoon, nucleotides 6:15 + 40:49) and group 2 (cyan cartoon, residues 16:39).

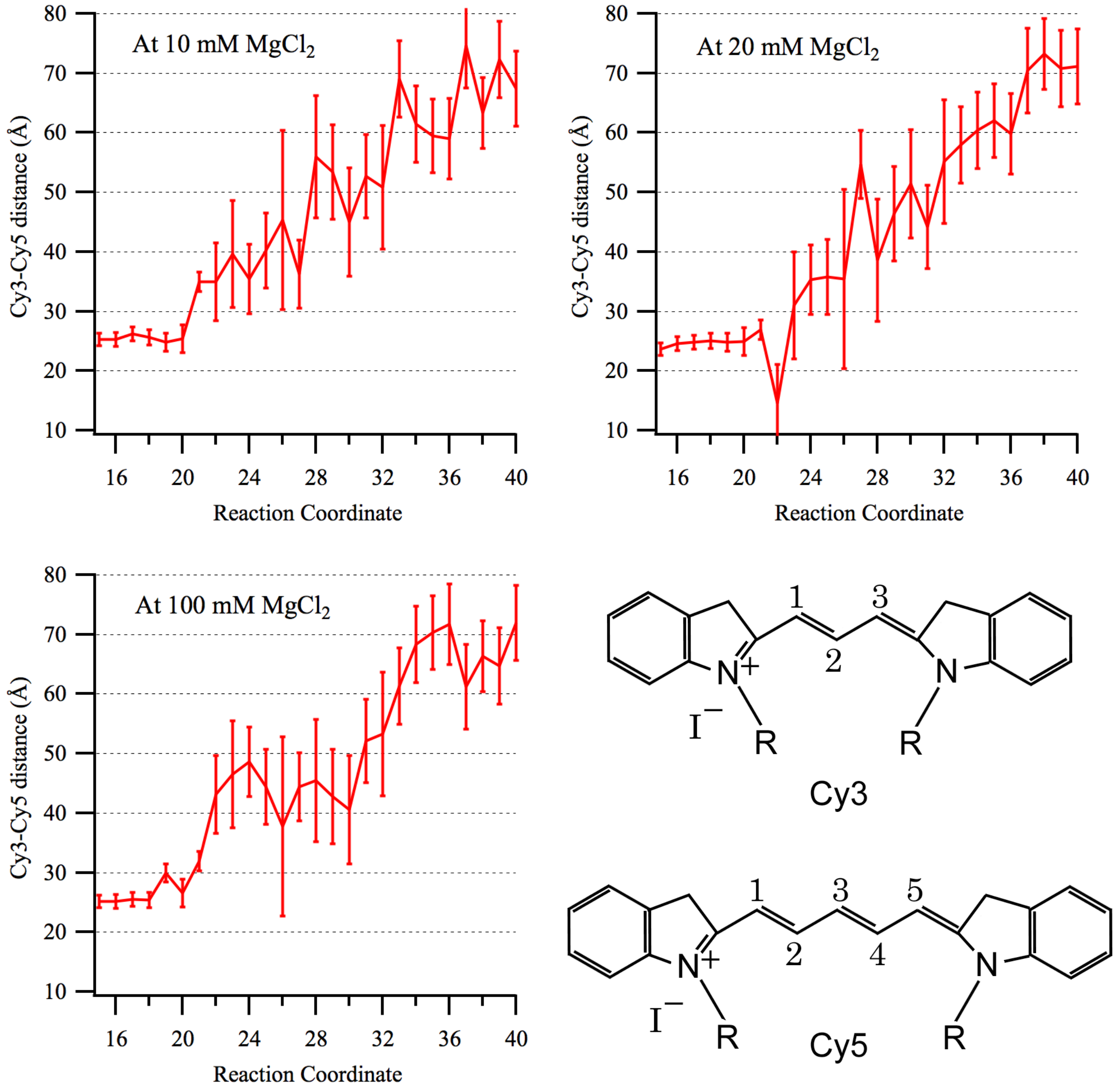


Figure S2. Interfluorophore distances based on the distance between C4' atoms of Ura 24 and Gua 54 along the reaction coordinate for Twister in 10, 20 and 100mM $MgCl_2$. Bottom right panel – Structures of Cy3 and Cy5 fluorophores.

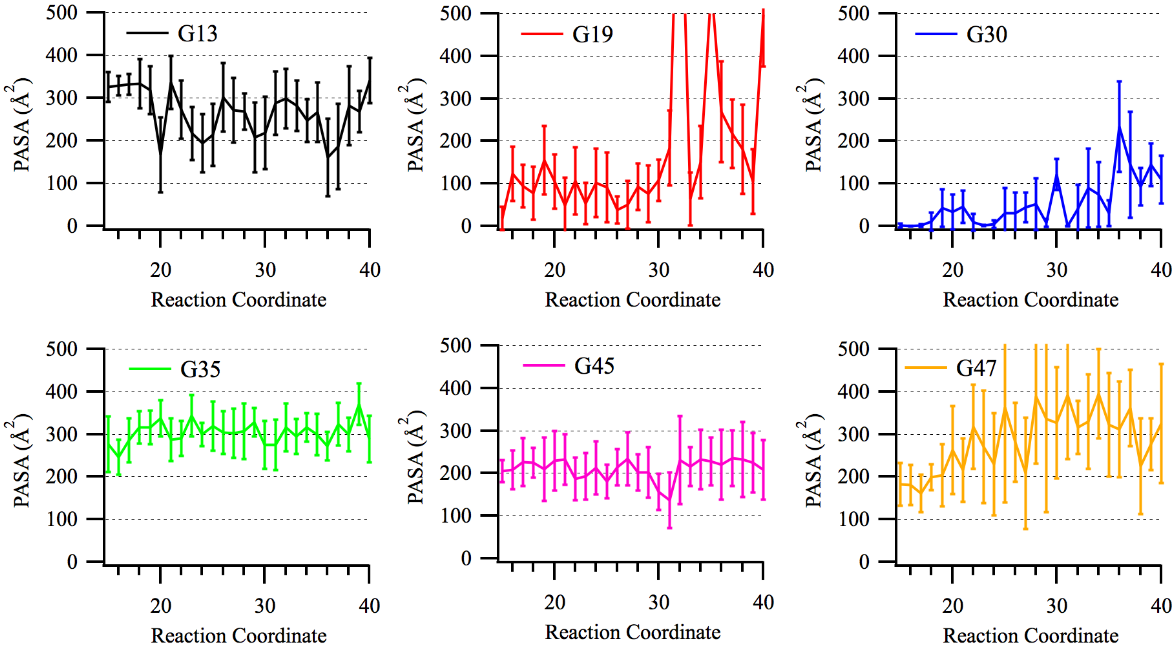


Figure S3: RNase T1 foot printing information. Protein accessible surface area (PASA) for the guanine nucleotides protected from cleavage by RNase T1 calculated at each window along the reaction coordinate with 0 mM $MgCl_2$. PASA is calculated as the solvent accessible surface area using a probe radius of 10 Å. The error bars represent one standard deviation.

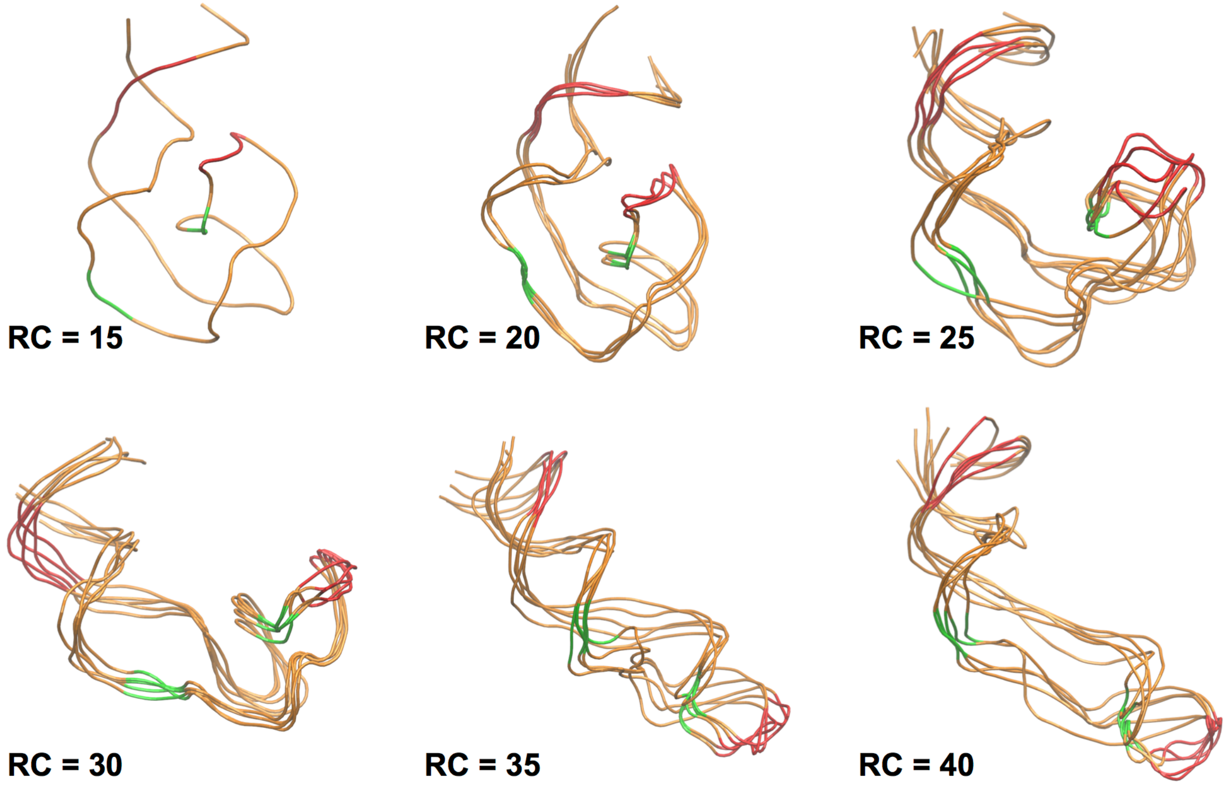


Figure S4. Center conformations of top 5 clusters from the 100mM $MgCl_2$ PMF at various values of the reaction coordinate. RMSD clustering analysis was performed with a cutoff of 2.5 Å from which the of top 5 clusters of conformations at each window were identified.

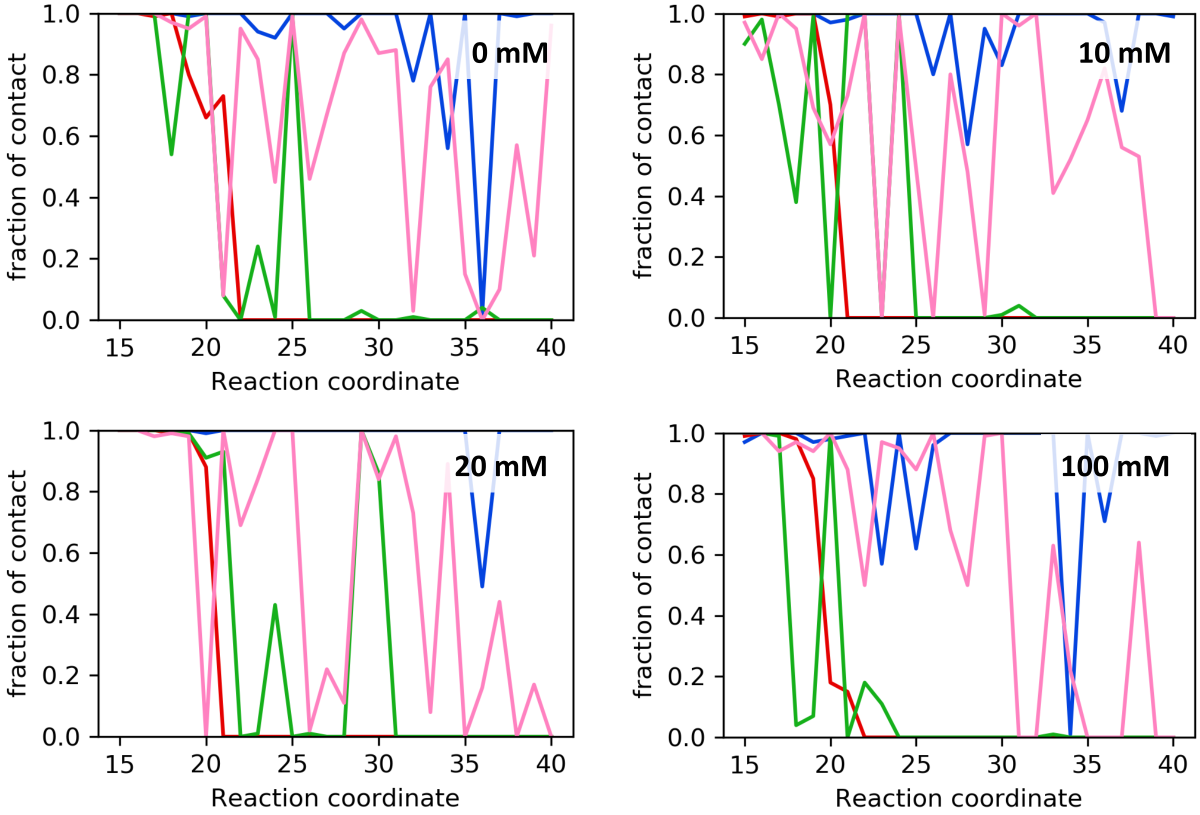


Figure S5. Fraction of tertiary contacts present along the reaction coordinate for the four different $MgCl_2$ concentrations. T1 – red, T2 – green, T3 – pink, T0 – blue.

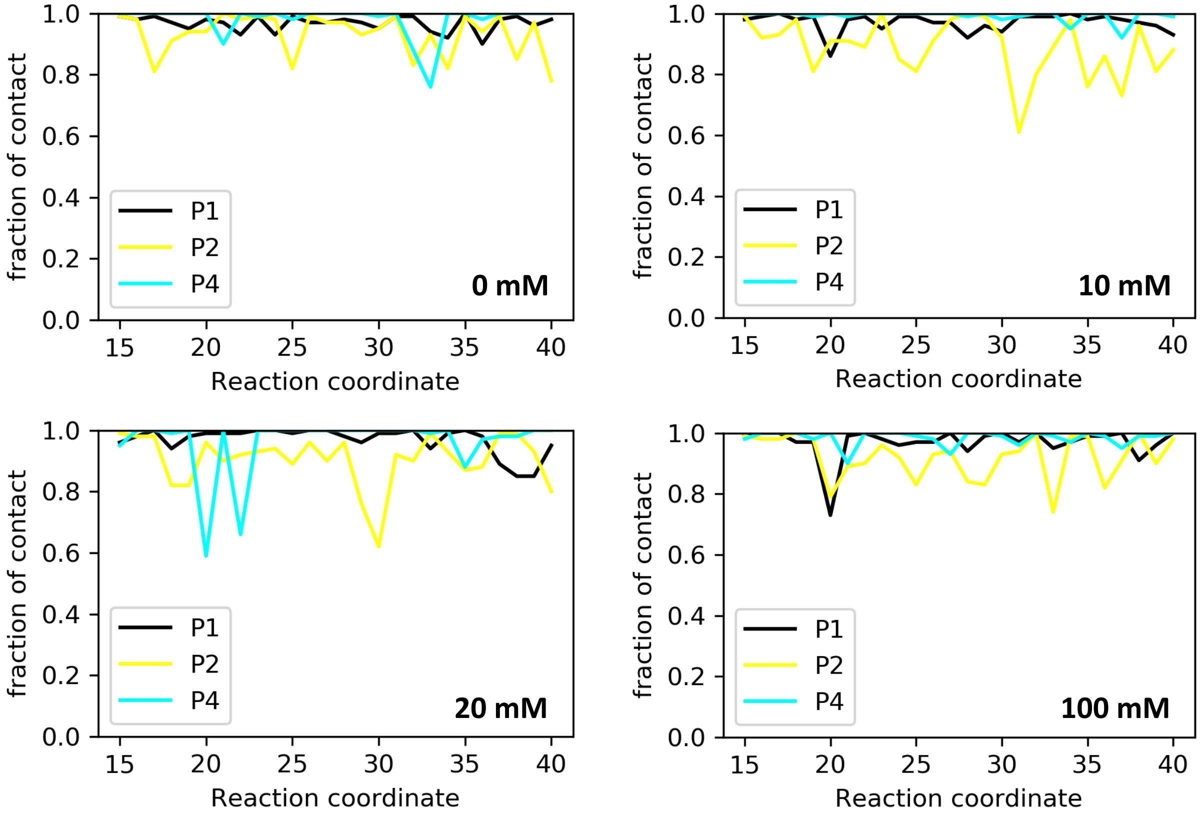


Figure S6. Fraction of secondary contacts present along the reaction coordinate for the four different $MgCl_2$ concentrations.

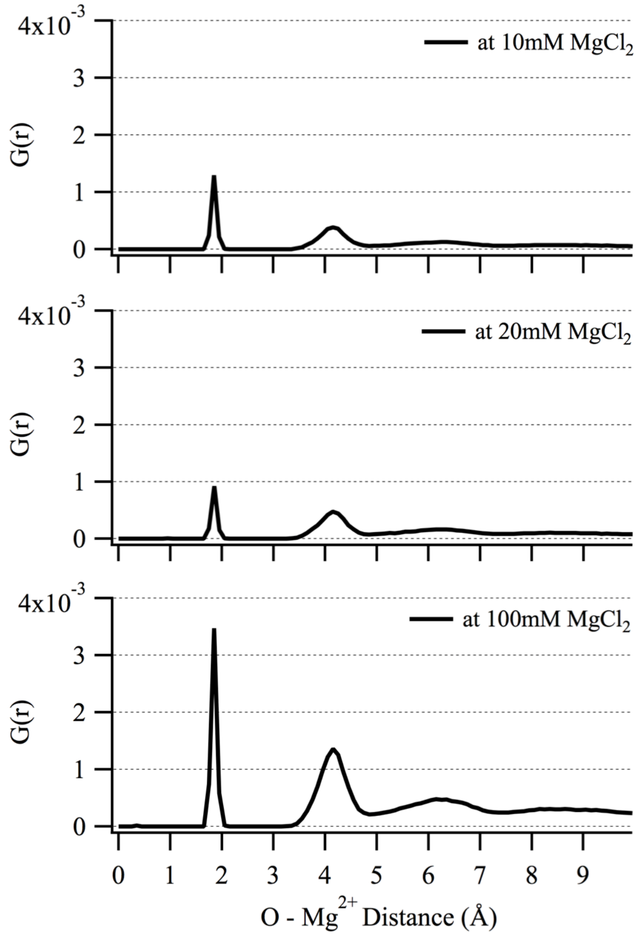


Figure S7. RDF of $Mg^{2+}$ ions around non-bridging phosphate oxygens (NBPO) in the 10, 20 and 100 mM $MgCl_2$ systems. A 9 Å distance between NBPOs was used to define those NBPO pairs that could be simultaneously interacting with $Mg^{2+}$ through outer shell interactions.

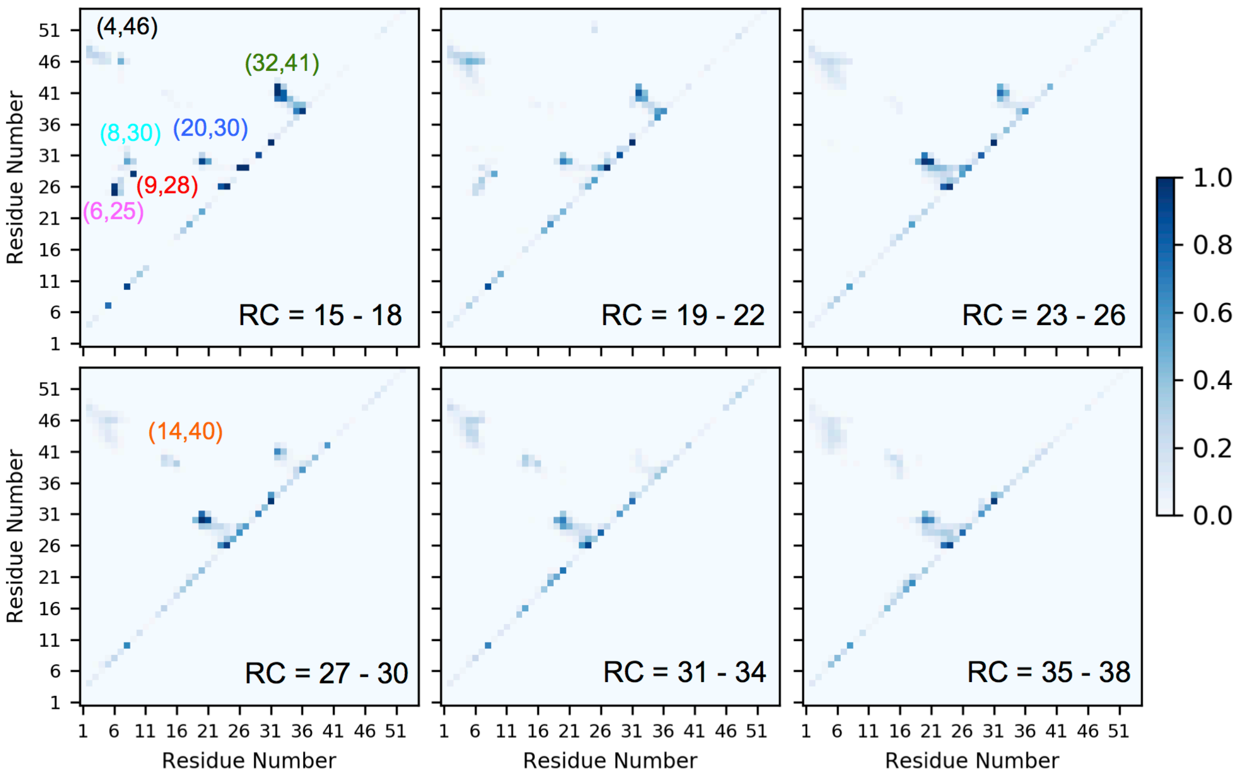


Figure S8. NBPO probability matrices from NBPO distance analysis for Twister from the GCMC-MD PMF at 0 mM $MgCl_2$. The probabilities were averaged over groups of 4 RC windows. RC = 15 – 18 corresponds to the fully folded structure.

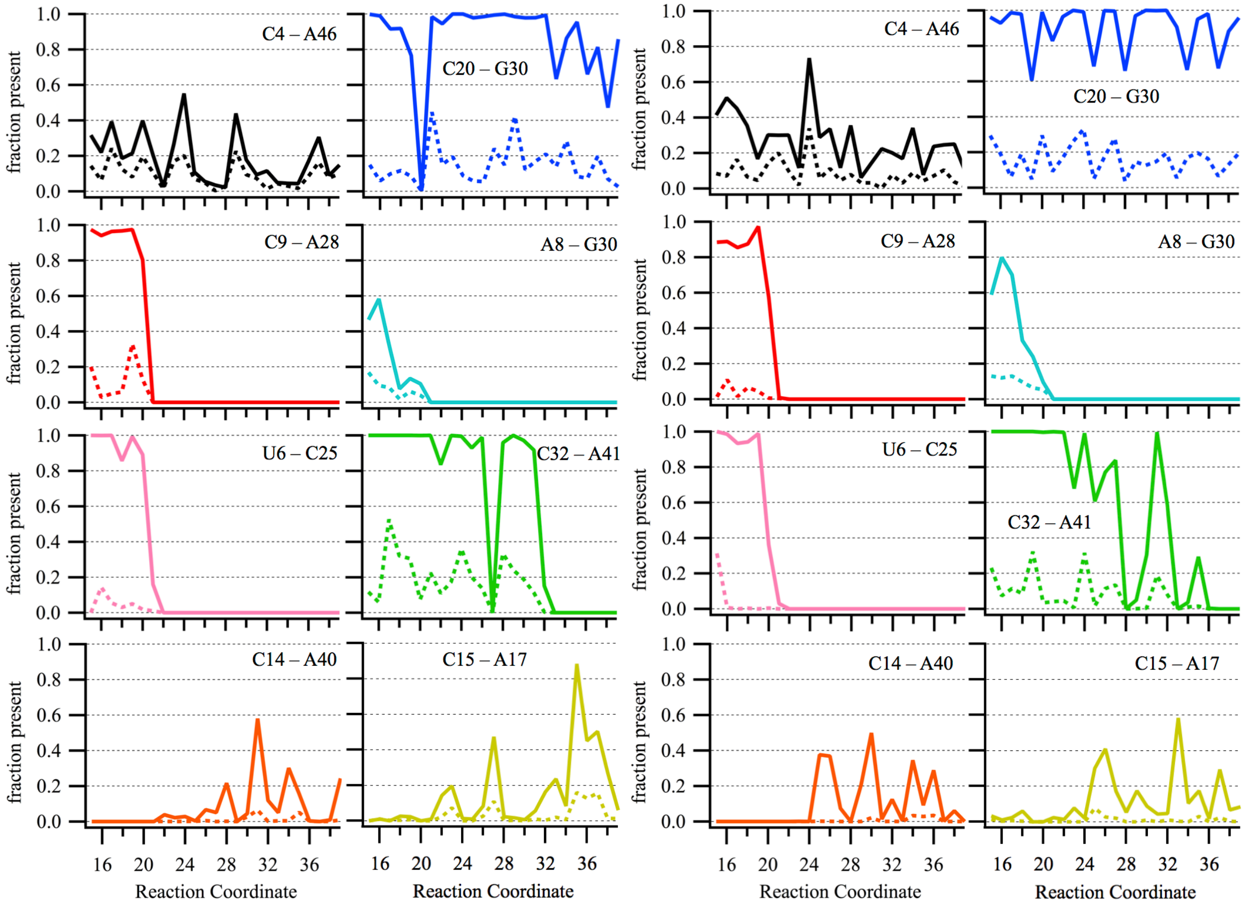


Figure S9. The probability of different pairs of nucleotides with NBPOs within 9 Å and the probability of a $Mg^{2+}$ ion coordinating the two NBPOs along the reaction coordinate. Left two panels – 20mM $MgCl_2$ Right two panels – 10mM $MgCl_2$

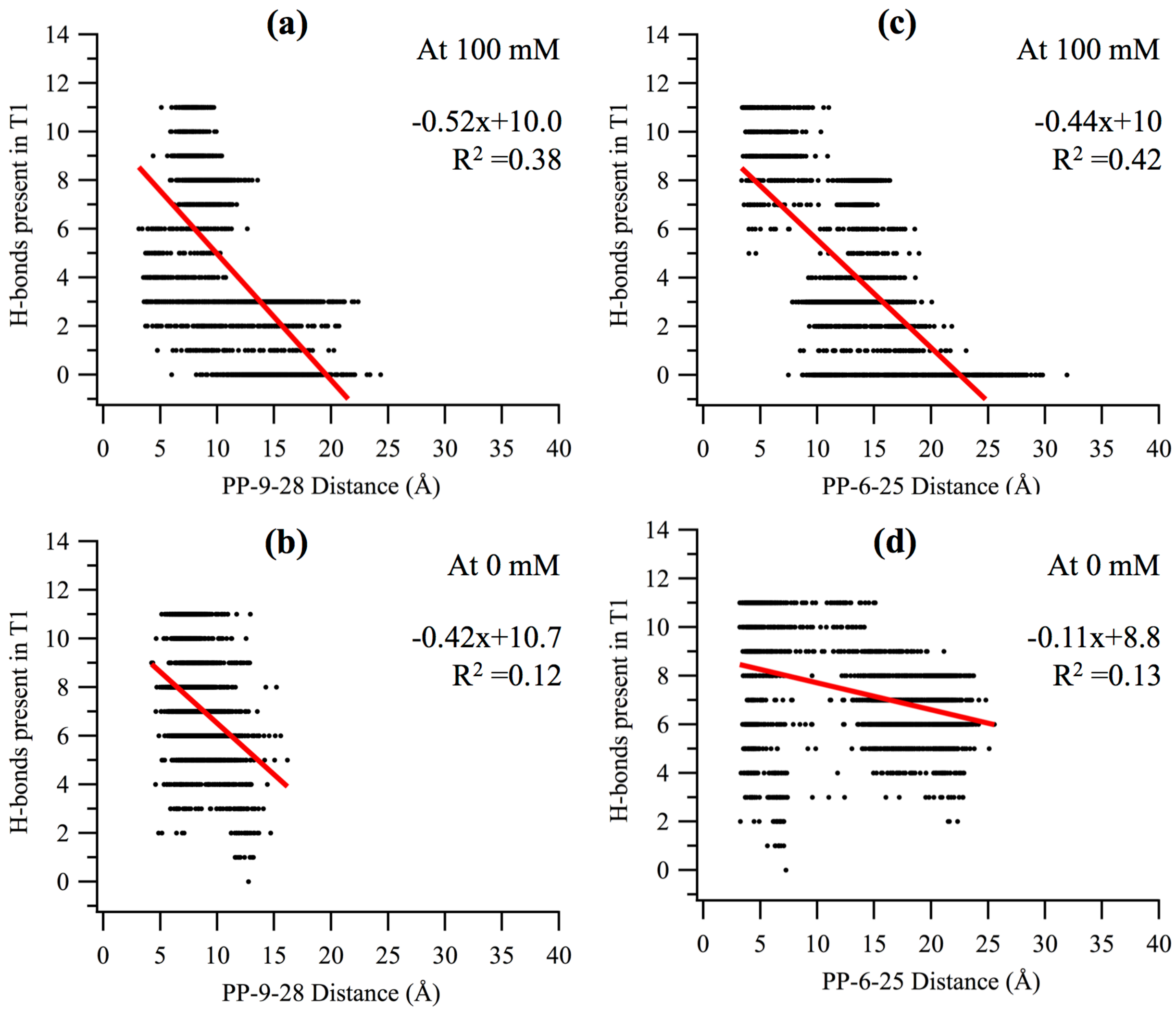


Figure S10) Correlation plots of the minimum NBPO distance versus the number of T1 tertiary contact WC hydrogen bonds for the pp-9-28 (a & b) and pp-6-25 (c & d) NBPO pairs. Data is from the RC = 19 to 21 windows of the PMF for the 100 mM and 0 mM $MgCl_2$ PMFs.

### SUPPORTING TABLES

Table S1a. Summed probability of selected regions of non-sequential NBPO pairs from the 0 and 100 mM $MgCl_2$ PMFs. Selected regions defining the NBPO pairs are listed in section 3 of the table. The values for the adjacent to the diagonal interactions (Figure 6, termed Near Diagonal) are based on the total summed probabilities (Table 2) minus the sum of the selected NBPO pair regions shown in this table.

| **100 mM** | | | | | | | | | |
|---|---|---|---|---|---|---|---|---|---|
| **RC** | **pp9-28** | **pp6-25** | **pp8-30** | **pp32-41** | **pp4-46** | **pp20-30** | **pp14-40** | **pp15-17** | **Near Diagonal** |
| 15 to 18 | 0.887 | 2.438 | 1.607 | 7.672 | 5.484 | 2.982 | 0.016 | 0.202 | 13.525 |
| 19 to 22 | 0.259 | 0.441 | 0.515 | 5.894 | 7.348 | 2.984 | 0.082 | 0.235 | 15.421 |
| 23 to 26 | 0.007 | 0.013 | 0.038 | 2.98 | 7.043 | 3.724 | 0.132 | 0.432 | 15.006 |
| 27 to 30 | 0 | 0 | 0.001 | 2.408 | 5.818 | 3.812 | 0.671 | 0.578 | 16.233 |
| 31 to 34 | 0 | 0 | 0 | 0.589 | 6.671 | 4.051 | 0.694 | 0.903 | 15.75 |
| 35 to 38 | 0 | 0 | 0 | 0.001 | 6.034 | 3.248 | 1.01 | 2.194 | 15.186 |
| **0 mM** | | | | | | | | | |
| **RC** | **pp9-28** | **pp6-25** | **pp8-30** | **pp32-41** | **pp4-46** | **pp20-30** | **pp14-40** | **pp15-17** | **Diagonal** |
| 15 to 18 | 0.962 | 2.833 | 1.409 | 6.914 | 2.49 | 2.055 | 0.144 | 0.092 | 12.963 |
| 19 to 22 | 0.52 | 1.147 | 0.335 | 4.569 | 4.613 | 2.033 | 0.082 | 0.254 | 14.525 |
| 23 to 26 | 0 | 0.001 | 0.014 | 3.634 | 3.397 | 4.897 | 0.248 | 0.559 | 15.752 |
| 27 to 30 | 0 | 0 | 0.001 | 2.155 | 2.276 | 4.524 | 1.073 | 0.541 | 15.123 |
| 31 to 34 | 0 | 0 | 0 | 0.695 | 3.832 | 3.655 | 1.419 | 0.661 | 15.155 |
| 35 to 38 | 0 | 0 | 0 | 0.004 | 2.988 | 3.944 | 0.988 | 1.095 | 16.144 |
| Regions defining the NBPO pairs | | | | | | | | | |
| pp9-28 = P(9,28)<br>pp6-25 = P(5:8,23:27)<br>pp8-30 = P(6:9,29,32)<br>pp32-41 = P(31:35,38:43) | | | | | pp4-46 = P(2:8,41:51)<br>pp20-30 = P(18:22,28,32)<br>pp14-40 = P(12:16,38:42)<br>pp15-17 = P(14:17,16:18) | | | | |

Table S1b) Standard deviations of difference between the summed probabilities at 100 and at 0 mM $MgCl_2$ for the individual non-sequential NBPO pairs for the results reported in Table 3.

| **RC** | **pp9-28** | **pp6-25** | **pp8-30** | **pp32-41** | **pp4-46** | **pp20-30** | **pp14-40** | **pp15-17** |
|---|---|---|---|---|---|---|---|---|
| 15 to 18 | 0.02 | 0.39 | 0.18 | 0.59 | 0.21 | 0.30 | 0.15 | 0.06 |
| 19 to 22 | 0.04 | 0.18 | 0.08 | 0.72 | 0.51 | 0.49 | 0.05 | 0.14 |
| 23 to 26 | 0.01 | 0.02 | 0.04 | 0.40 | 0.90 | 0.40 | 0.16 | 0.18 |
| 27 to 30 | 0.00 | 0.00 | 0.00 | 1.13 | 0.68 | 0.36 | 0.27 | 0.13 |

| | | | | | | | | |
|---|---|---|---|---|---|---|---|---|
| 31 to 34 | 0.00 | 0.00 | 0.00 | 0.27 | 1.29 | 0.30 | 0.20 | 0.24 |
| 35 to 38 | 0.00 | 0.00 | 0.00 | 0.00 | 1.12 | 0.37 | 0.73 | 0.14 |

Table S1c) Difference between the summed probabilities at 100 and at 0 mM $MgCl_2$ for the individual non-sequential NBPO pairs and for the non-sequential pairs adjacent to the diagonal on Figure 6 over RC = 19 to 22 Å. The RC=22 window was excluded as the number of T1 hydrogen bonds was zero; exclusion of the RC=22 window led to the difference between the 100 and 0 mM 9-28 and 6-25 probabilities being -0.35 and -0.95, respectively, which are of larger magnitude than the values of -0.26 and -0.71 presented in Table 3 for RC = 19 to 22.

| RC=19 to 21 | Total | pp9-28 | pp6-25 | pp8-30 | pp32-41 | pp4-46 | pp20-30 | pp14-40 | pp15-17 | Near Diagonal |
|---|---|---|---|---|---|---|---|---|---|---|
| At 100mM | 33.55 | 0.34 | 0.58 | 0.61 | 6.76 | 7.07 | 2.79 | 0.10 | 0.20 | 15.09 |
| At 0mM | 29.40 | 0.69 | 1.53 | 0.42 | 5.79 | 5.28 | 1.18 | 0.01 | 0.18 | 14.32 |
| Diff (100-0) | 4.16 | -0.35 | -0.95 | 0.19 | 0.97 | 1.79 | 1.62 | 0.09 | 0.02 | 0.77 |

Table S2. Hydration free energies of the ions and water and the minimum and maximum values of $\mu_{ex}$ used in the PME/GCMC protocol.

| Fragment | Hydration FE (kcal/mol) | $\mu_{ex}^{min}$ (kcal/mol) | $\mu_{ex}^{max}$ (kcal/mol) |
|---|---|---|---|
| $Mg^{2+}$ | -437.38 | -837.38 | -37.38 |
| $K^{+}$ | -70.51 | -170.51 | 30.51 |
| $Cl^{-}$ | -81.26 | -81.26 | -81.26 |
| Water | -5.60 | -5.60 | -5.60 |

### Supporting Text 1. Validation of the PME GCMC-MD protocol

To validate the oscillating $\mu_{ex}$ GCMC $Mg^{2+}$ ion sampling protocol we calculated the $Mg^{2+}$ ion distribution in four RNA structures, VS ribozyme (*PDB 2MIS*) (1), BWYV pseudoknot (*PDB 1L2X*) (2), Twister ribozyme from *O. Sativa* (*PDB 4OJI*) (3), and preQ1 riboswitch (*PDB 3FU2*) (4). The former two were previously analyzed by our group for characterization of $Mg^{2+}$ distributions around native conformations using a similar oscillating $\mu_{ex}$ GCMC-MD based approach (5). However, that study applied a non-bond truncation scheme for the electrostatic interactions. In the present work, a recently published extension of the oscillating $\mu_{ex}$ GCMC method (6) that includes particle mesh Ewald (PME) (7) for the treatment of long-range electrostatics was applied in conjunction with a new $\mu_{ex}$ oscillation scheme for the excess chemical potential to sample the interactions of $Mg^{2+}$ with RNA. To prepare the systems for validation of the new PME GCMC-MD protocol, all divalent ions and waters present in the crystal structures were removed. Other monovalent ions ($Na^{+}$ and $K^{+}$) were retained in their experimental positions. For the VS ribozyme stem-loop VI solution NMR structure, the first model (out of 20 in the NMR ensemble) was used following the protocol from our previous study (5). For Twister ribozyme, two missing nucleotides in the crystal structure were added by using the internal coordinates in CHARMM (8). A stepwise restrained minimization of the coordinates to relax the local conformation of the two nucleotides. For the preQ1 riboswitch the PRF ligand was removed. All four RNAs were solvated in a 65 Å cubic waterbox with 50 mM $MgCl_2$ and 100 mM KCl. The average size of the systems was ~31500 atoms. The systems were minimized and equilibrated with restraints on the backbone atoms using CHARMM (8). The same conditions were used for the equilibration MD simulations as described in the methods. The final conformations were used to start the GCMC-MD validation runs where the backbone atoms were harmonically restrained during the MD part to preserve the native conformation. The validation runs involved 4 replicates of 50 cycles with 5 ns MD run per cycle. In total 1000 ns of MD simulation was performed for each RNA.

The validation simulations on the 4 RNAs show that the PME-based GCMC approach for sampling the ions around the RNA is more effective than the previous protocol (5). Analysis involved the distribution of $Mg^{2+}$ ions over all 2000 frames from the 4 replicates × 50 cycles performed on each of the 4 systems. Following alignment of the frames with respect to the RNA backbone atoms of the experimental structures, clustering of the $Mg^{2+}$ ion positions around the RNA was performed based on RMSD of ion positions. Figure T1 shows the clusters of $Mg^{2+}$ ions colored with respect to their cluster size. Compared to the respective X-ray or NMR structures (1-4), almost all the experimental $Mg^{2+}$ binding sites have been successfully sampled and represented by clusters with significant number of members. Table T1 shows the distance to the nearest cluster for crystallographic positions of $Mg^{2+}$ for the Twister ribozyme. In addition, the $Mg^{2+}$ positions from the GCMC-MD runs with Twister are well distributed with respect to inner-shell, outer-shell and weakly bound positions (Figure T2). This indicates that the PME GCMC-MD protocol is capable and efficient in finding binding sites associated with a distribution of $Mg^{2+}$ coordination types.

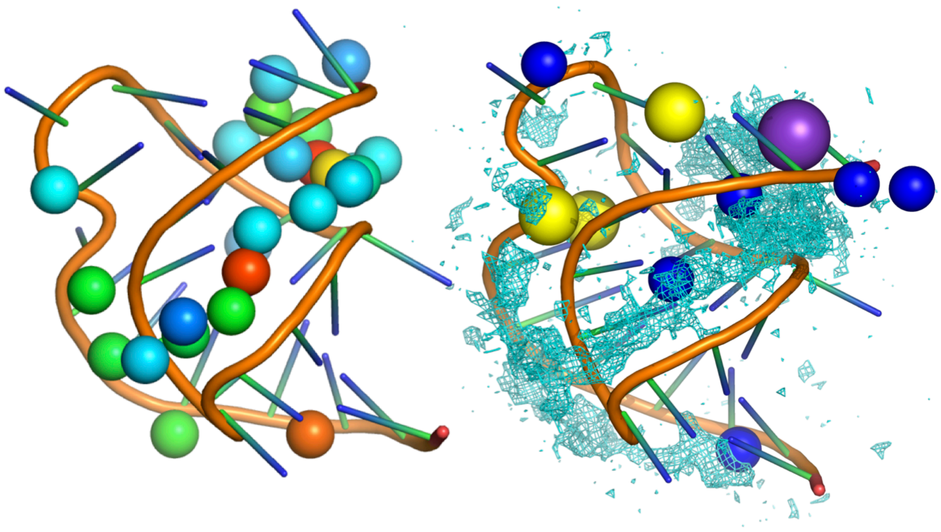

BWYV Pseudoknot

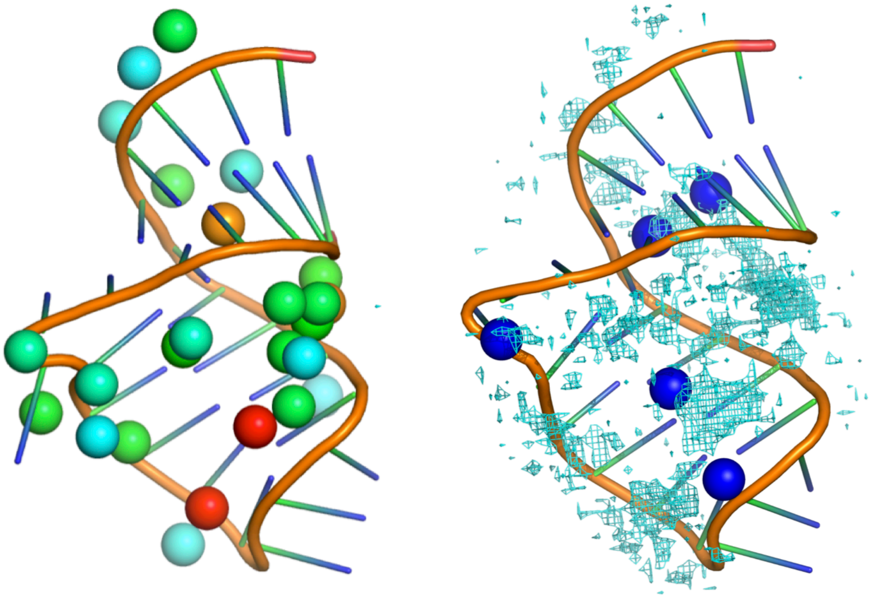

VS Ribozyme

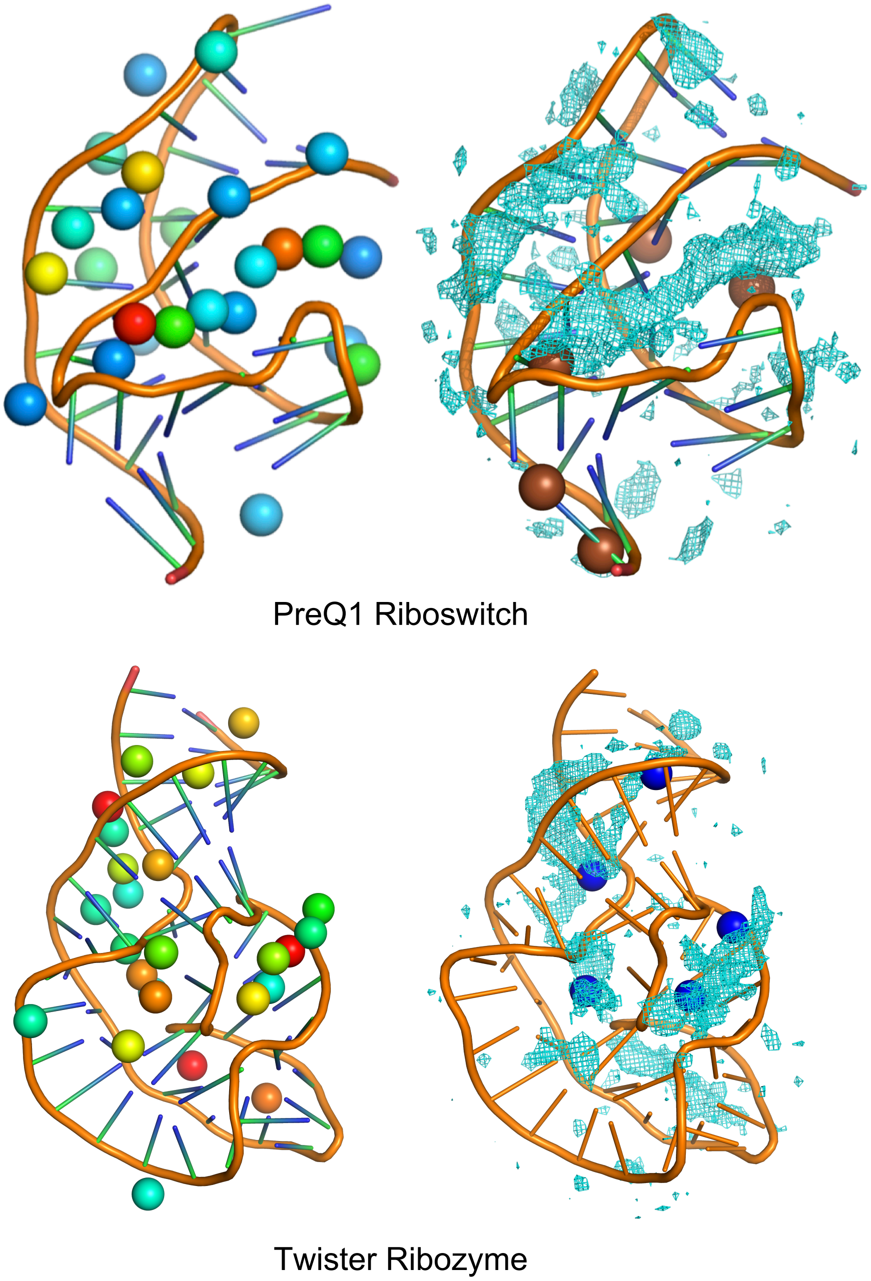


Figure T1. The left panels show centers of top 25 clusters calculated from the $Mg^{2+}$ ion distribution from the GCMC-MD simulations around the 4 RNA systems. On the left panels, spheres representing the cluster centers are colored from red to blue. The right panels illustrate the binding sites of ions captured in X-ray or NMR experiments (1-4). $Mg^{2+}$ occupancy maps with GFE cutoff of -2 kcal/mol are shown. On the right panels, Mg - Dark blue spheres, Na - Yellow spheres, K - Purple sphere, Ca - brown spheres.

Table T1. Minimum distance between experimental $Mg^{2+}$ binding sites for Twister and cluster center for $Mg^{2+}$ positions sampled in the 100 mM $MgCl_2$ GCMC/MD validation runs.

| Crystal structure identifier for Mg ion | Distance to nearest cluster center (Å) |
|---|---|
| MG 101 | 3.16 |
| MG 102 | 5.61 |
| MG 103 | 2.02 |
| MG 104 | 3.22 |
| MG 105 | 2.10 |

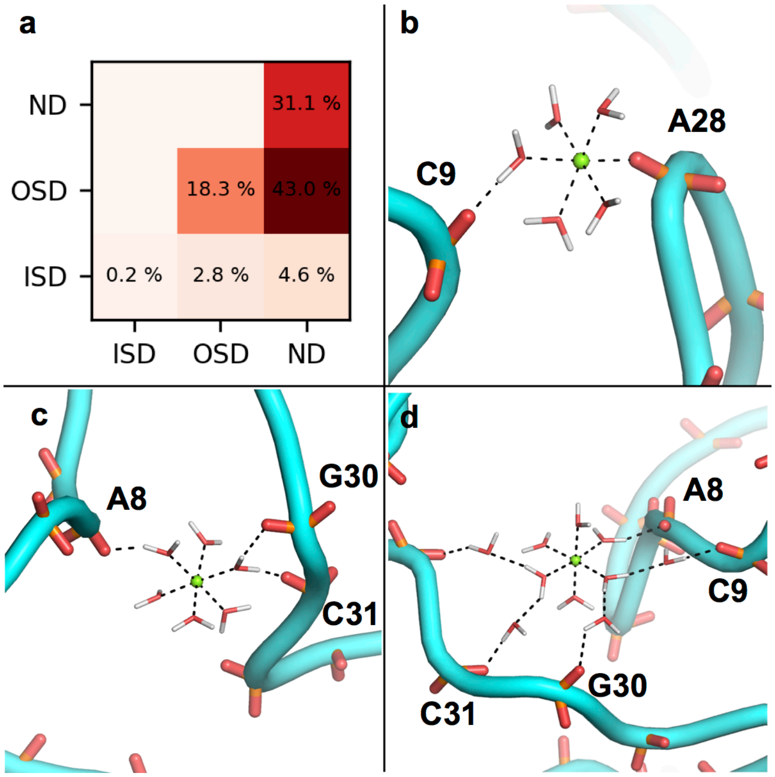


Figure T2. a) Distribution of combination of interactions between $Mg^{2+}$ and NBPOs over all the simulations. Interaction types are direct-interaction (Inner-Shell-Dehydrated: ISD), indirect-interaction (Outer-Shell-Dehydrated: OSD) and diffused-interaction (Non-Dehydrated: ND). 70% $Mg^{2+}$ ions were found to be participating in at least one strong interaction, either ISD or OSD, for one of the NBPOs in each pair. Very rarely both NBPOs are simultaneously coordinated at inner-shell level, while OSD+ND or ND+ND types of combination occur more frequently. Examples of the types of combinations include b) ISD with OSD interactions providing strong coordination with phosphate groups c) OSD with OSD interactions with phosphate groups and d) OSD with ND interactions with phosphate groups.
